# Computational Mechanisms of Neuroimaging Biomarkers Uncovered by Multicenter Resting-State fMRI Connectivity Variation Profile

**DOI:** 10.1101/2024.04.01.587535

**Authors:** Okito Yamashita, Ayumu Yamashita, Yuji Takahara, Yuki Sakai, Yasumasa Okamoto, Go Okada, Masahiro Takamura, Motoaki Nakamura, Takashi Itahashi, Takashi Hanakawa, Hiroki Togo, Yujiro Yoshihara, Toshiya Murai, Tomohisa Okada, Jin Narumoto, Hidehiko Takahashi, Haruto Takagishi, Koichi Hosomi, Kiyoto Kasai, Naohiro Okada, Osamu Abe, Hiroshi Imamizu, Takuya Hayashi, Shinsuke Koike, Saori C. Tanaka, Mitsuo Kawato, Brain/MINDS Beyond Human Brain MRI Group

## Abstract

Resting-state functional connectivity (rsFC) is increasingly used to develop biomarkers for psychiatric disorders. Despite progress, development of the reliable and practical FC biomarker remains an unmet goal, particularly one that is clinically predictive at the individual level with generalizability, robustness, and accuracy. In this study, we propose a new approach to profile each connectivity from diverse perspective, encompassing not only disorder-related differences but also disorder-unrelated variations attributed to individual difference, within-subject across-runs, imaging protocol, and scanner factors. By leveraging over 1500 runs of 10-minute resting-state data from 84 traveling-subjects across 29 sites and 900 participants of the case-control study with three psychiatric disorders, the disorder-related and disorder-unrelated FC variations were estimated for each individual FC. Using the FC profile information, we evaluated the effects of the disorder-related and disorder-unrelated variations on the output of the multi-connectivity biomarker trained with ensemble sparse classifiers and generalizable to the multicenter data. Our analysis revealed hierarchical variations in individual functional connectivity, ranging from within-subject across-run variations, individual differences, disease effects, inter-scanner discrepancies, and protocol differences, which were drastically inverted by the sparse machine-learning algorithm. We found this inversion mainly attributed to suppression of both individual difference and within-subject across-runs variations relative to the disorder-related difference by weighted-averaging of the selected FCs and ensemble computing. This comprehensive approach will provide an analytical tool to delineate future directions for developing reliable individual-level biomarkers.

## Introduction

Mental disorders have become a serious social problem in recent years. Epidemiological and economic analyses have indicated that their global impact is substantial in terms of human health and social welfare^1^. However, current diagnostic methods, which are based on self-reported symptoms or those identified during medical interviews, are insufficient for treatment optimization. Biomarkers based on genes, blood analyses, and neuroimaging data could overcome these limitations^2^.

Resting-state functional connectivity (rsFC) is one of the most promising approaches for developing psychiatric disorder biomarkers^3^. This method assesses brain functional network by quantifying coactivation of spontaneous fluctuations across brain regions using a noninvasive brain measurement technique, usually functional magnetic resonance imaging (fMRI)^4,5^. Extensive research has highlighted the relevance of functional connectivity (FC) in relation to individual characteristics^6–9^, task activities^10,11^, brain states^12^, anatomical structure^13^ and neuronal signals^14–16^. Owing to its simplicity, versatility, interpretability, and sensitivity to individual variations, the FC biomarker shows great promise for objective diagnosis^17–20^, personalized treatment selection^21,22^, and neuromodulation target identification in psychiatry^23–26^.

Despite progress, development of the reliable and practical FC biomarker remains an unmet goal, particularly one that is clinically predictive at the individual level with generalizability, robustness, and accuracy^27–29^. Although altered functional connections between patient groups and healthy controls have been identified^30–33^, the individual-level classifications can be only achieved with the help of the machine-learning algortihms. In our multicenter study, we successfully developed major depressive disorder (MDD), schizophrenia (SCZ), and autism spectrum disorder (ASD) biomarkers using ensemble sparse classifiers, which generalized well across data from various centers^18–20^ and maintained consistent performance on new data (anterograde generalization)^34^. However, its discrimination ability evaluated with completely independent datasets, with areas under the curve of 0.74, 0.82, and 0.66–0.81 for MDD, SCZ, and ASD, respectively, may not yet meet high standards.

Two key obstacles must be overcome to develop a reliable and practical biomarker. First, there is the issue of limited dataset sizes used in machine-learning training, which is problematic given patient heterogeneity. Despite recognizing the value of multicenter rsFC studies for gathering large-scale data and creating biomarkers practical with real-world data^28,35–37^, there is still an incomplete comprehensive and quantitative understanding of the variability in FC across multiple centers. Second, we face a knowledge gap regarding how machine-learning algorithms facilitate individual classification and what factors limit achieving higher accuracy.

This study aimed to comprehensively and quantitatively evaluate the effects of various factors on FC and machine-learning–based biomarker outputs to delineate future directions for developing reliable individual-level biomarkers. We leveraged data from two major decade-spanning Japanese projects, the Strategic Research Program for Brain Sciences (SRPBS) (2012–2018) and Brain/Minds Beyond (BMB) (2018–2024). These projects have been uniquely featured for their extensive data with the collection from numerous traveling-subjects (84 participants) and approximately 10,000 participants including several thousand patients with diverse psychiatric and neurological disorders across multiple centers^37,38^. This study analyzed approximately 2,400 runs of 10-minute eyes-open resting-state fMRI data from the BMB and SRPBS traveling-subject datasets, as well as an SRPBS multi-disorder dataset (Fig. 1). We found hierarchical variations in individual FC, ranging from run-to-run variations, individual differences, disease effects, inter-scanner discrepancies, and protocol differences. The sparse machine-learning algorithm proposed in our previous study^19^ can effectively prioritize disease effects via optimal selection of FCs, their weighting and ensemble averaging, and drastically inverted the above order of variability factors. More specifically, we revealed three distinct computational mechanisms that improve our biomarker’s signal to noise ratio (disorder effect/participant related variabilities) almost 15 times. These findings render our rsFC biomarkers practical for clinical applications and highlight the need to further minimize individual differences and within-subject run-to-run variability to improve predictive modeling.

**Figure 1.**
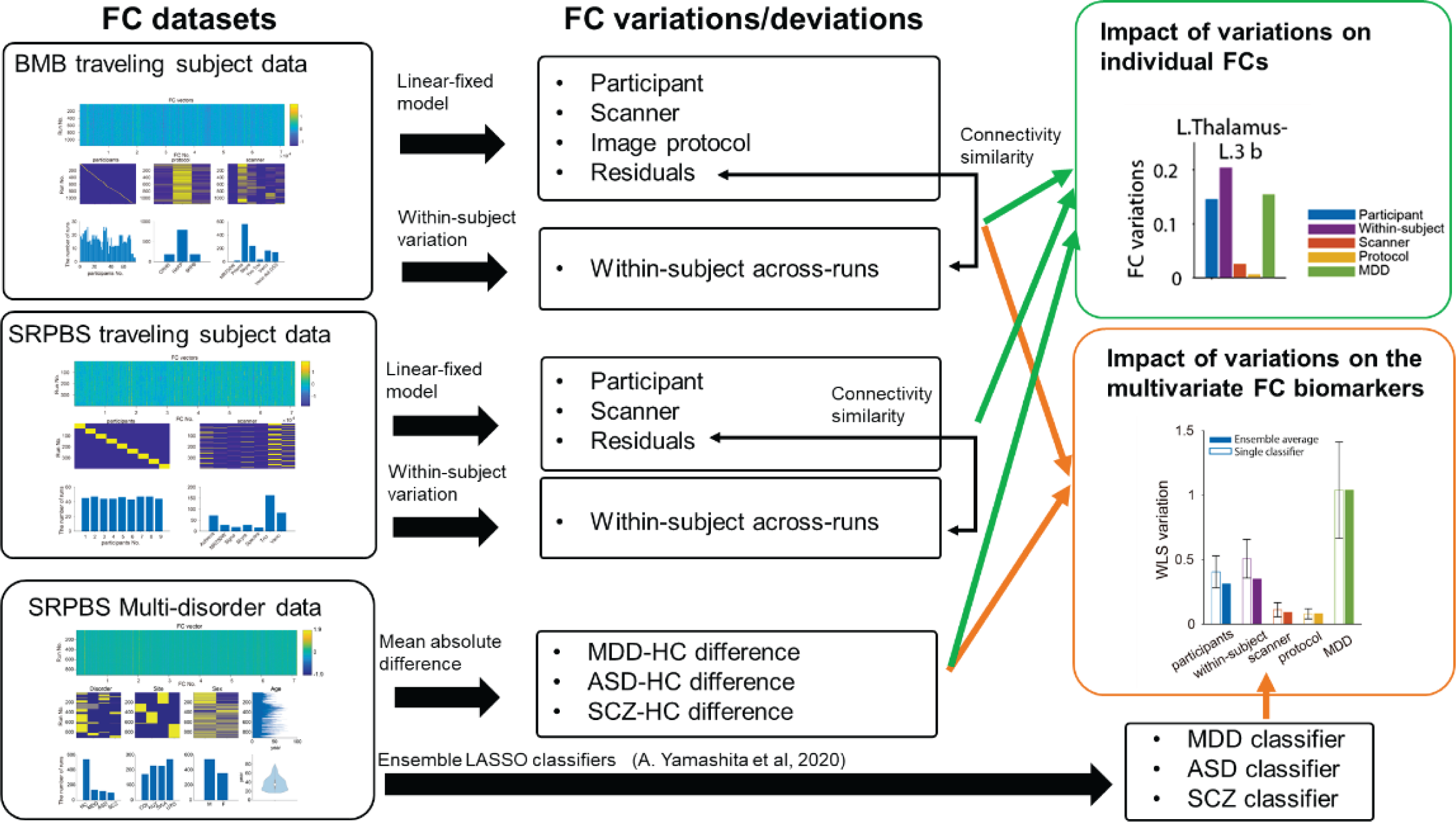
Schematic of the data analysis protocol used in this study. Two traveling-subject datasets from two nationwide projects conducted in Japan (Brain/Minds Beyond (BMB) and Strategic Research Program for Brain Sciences (SRPBS), along with a multi-disorder dataset from SRPBS, were analyzed in the present study to facilitate a more comprehensive understanding of functional connectivity (FC) variations related to individual differences, scanner differences, imaging protocol differences, and unexplained residual components in comparison to disorder-related FC variations. First, we computed the FC variations due to each factor and residual components using a carefully designed linear fixed effects model. We separately computed the within-subject across-runs FC variations using a subset of the traveling-subject datasets (inclusion criteria: subjects with data from at least six runs under a single measurement condition). The FC was characterized by the magnitude of the FC variations attributed to differences in individuals (participants), scanners, imaging protocols, within-subject across-run differences, and disorder differences. Furthermore, we evaluated the impact of each factor’s FC variations on the outcomes of the multivariate FC biomarker we previously developed using least absolute shrinkage and selection operator (LASSO) ensemble classifiers^19^.

## Results

### FC variation analysis in the BMB traveling-subject dataset

We applied a three-factor linear fixed effects model to the BMB traveling-subject dataset to investigate FC variations due to participant, imaging protocol, and scanner factors and unexplained residual components for each connection.

The magnitude distributions of the FC variations across all connections (71,631 connections using Glasser’s Multimodal Parcellation (MMP) atlas) are presented in Fig. 2(a). The median values (5^th^–95^th^ percentile) for the participant, protocol, and scanner factors were 0.107 (0.066–0.192), 0.016 (0.004–0.042), and 0.0259 (0.012–0.055), respectively; the magnitude of the median value of the unexplained residuals was larger than that of the three factors (0.160 (0.146–0.183)). The distribution for the the participant factor (individual differences) was broad, whereas the distributions for the protocol, scanner, and residual factors were narrower.

**Figure 2.**
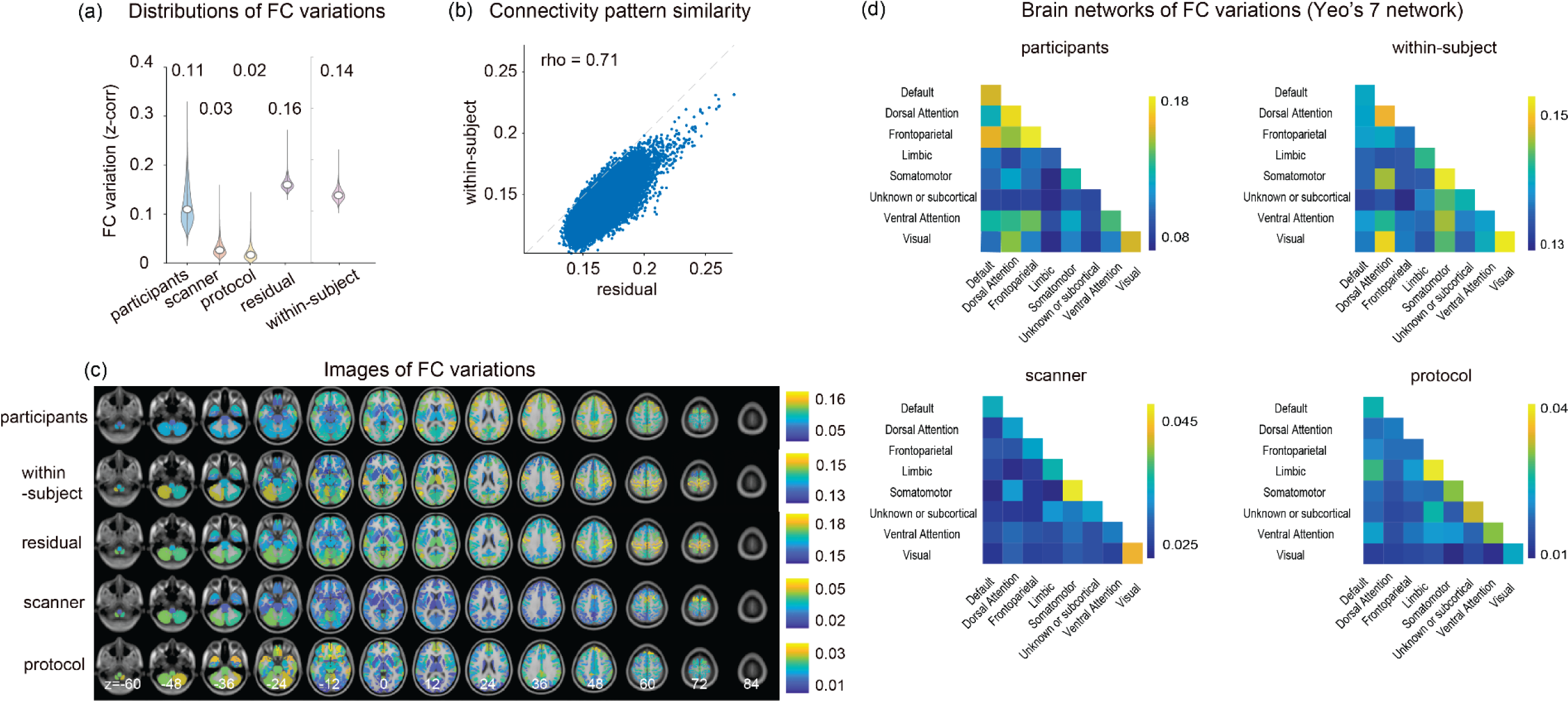
Analysis of functional connectivity (FC) variations for the Brains/Minds Beyond (BMB) traveling-subject dataset based on Glasser’s Multimodal Parcellation (MMP) atlas. A linear fixed effects model with three factors was applied to the BMB traveling-subject dataset to investigate the FC variations due to the participant (73 subjects), imaging protocol (three protocols), and scanner (six scanners) factors, and the unmodeled residual component, which was characterized on the basis of separately computed within-subject FC variations. (a) Distributions of the magnitudes of the FC variations due to the participant, scanner, protocol, residual component, and within-subject factors. Each violin plot summarizes the whole-brain FC variations across 71,631 connections. The median value of each distribution is shown above each violin plot. (b) Comparison of the connectivity pattern similarity between the residual component and within-subject variations. Each dot corresponds to one connectivity. (c) Brain mapping of the FC variations due to the participant, within-subject, residual, scanner, and imaging protocol factors. (d) Brain networks of the FC variations summarized using Yeo’s 7-network parcellation. All reported values are represented by Fisher’s z-transformed Pearson correlation coefficients.

To elucidate the origin of the FC variation attributed to unexplained residuals, we investigated associations between it and within-subject across-run variations on the basis of connectivity pattern similarity (i.e., the rank correlation between residual and within-subject FC variation patterns). We observed a strong association between the residual and within-subject FC variations (correlation coefficient: 0.71) (Fig. 2(b)). The median magnitude of the within-subject FC variation was slightly smaller than that of the residuals (0.138 vs 0.160, respectively), suggesting another unknown factor was contributing to the residual components (Fig. 2(a)). This strong association (rank correlation=0.57) was observed even when the data used to calculate the residual component (595 runs, 42 participants) and within-subject (201 runs, 31 participants) variations were completely separated (Supplementary Fig. 1), indicating a substantial portion of the residual component effects could be accounted for by the within-subject FC variation.

Subsequently, we examined brain regions and networks affected by each factor, generating maps of the FC variations due to participant, protocol, and scanner factors and within-subject variations and residual components. The imaging value for a specific brain region was calculated as the average amplitude across all connections linking that specific brain region to all others. The participant factor variation was large in regions associated with the dorsal attention, frontoparietal, and default mode networks. The within-subject variation was large over the whole brain, with larger variations in the somatosensory, motor, and visual cortices, and certain dorsal attention network regions. Large protocol-related variation was observed in the anterior and inferior parts of the brain, including the orbitofrontal cortex, gyrus rectus, and olfactory regions. Large scanner-related variation was observed in the superior frontal gyrus and cerebellum at the top and bottom of the brain, respectively.

To characterize differences between two imaging protocols or scanners in the FC space, we defined the distance between them as the mean absolute difference between the corresponding estimated parameter vectors over all connections. We observed large distances between the Harmonized Protocol (HARP) and SRPBS protocols and between the SRPBS and Connectomes Related to Human Diseases (CRHD) protocols compared with the distances between the HARP and CRHD protocols, as expected, as well as large distances between the MR750W scanner from General Electric (GE) and the remaining Siemens scanners (Supplementary Fig. 2(a)(b)).

### FC variation analysis of the SRPBS traveling-subject dataset

To assess commonalities and differences in the aforementioned findings, the same analyses were applied to the SRPBS traveling-subject dataset using two-factor linear fixed effects modeling with participant and scanner factors.

The distributions of the FC variation magnitudes across all connections (71,631 connections from Glasser’s MMP atlas) and the association between residual component and within-subject FC variations are presented in Fig. 3(a), (b). The median values and quantiles of the participant and scanner factors were 0.080 (0.038–0.158) and 0.037 (0.019–0.071), respectively. The largest magnitude was observed for the residual components (0.156 (0.138–0.189)). The distribution for the participant factor was broader than that of the other factors. High connectivity pattern similarity was observed between the residual and within-subject variations (correlation coefficient: 0.69), with the median magnitude of the latter being smaller than that of the former (0.133 vs 0.160, respectively). These results were consistent with those in the BMB traveling-subject dataset analysis (Fig. 2(a), (b)).

**Figure 3.**
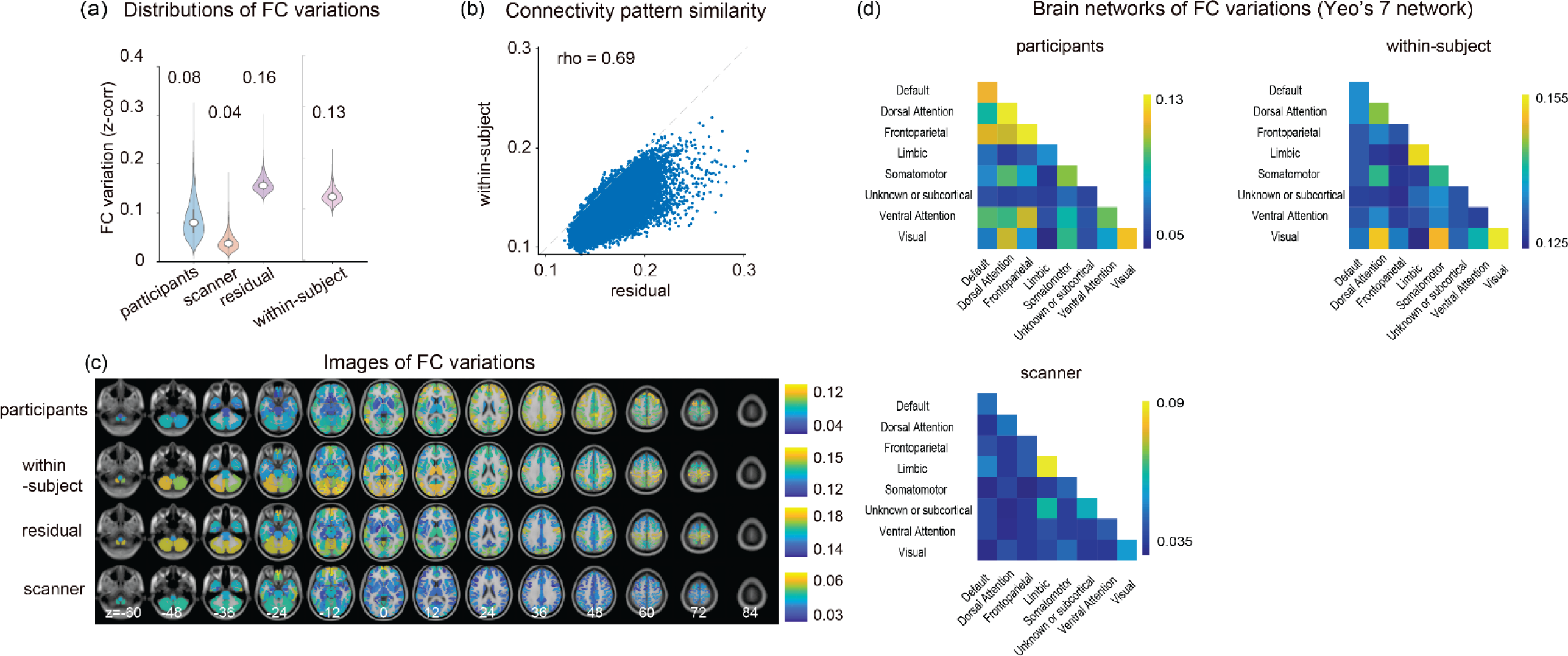
Analysis of functional connectivity (FC) variations for the Strategic Research Program for Brain Sciences (SRPBS) traveling-subject dataset based on Glasser’s Multimodal Parcellation (MMP) atlas. A linear fixed effects model with two factors was applied to the SRPBS traveling-subject dataset to investigate the FC variations due to the participant (nine subjects) and scanner (seven scanners) factors and the unmodeled residual component. The unmodeled residual component was characterized by separately computed within-subject FC variations. (a) The distributions of the magnitudes of the FC variations due to the participant, scanner, residual component, and within-subject factors. Each violin plot summarizes the whole-brain FC variations across 71,631 connections. The median value of each distribution is shown above each violin plot. (b) Comparison of the connectivity pattern similarity between the residual component and within-subject variations. Each dot corresponds to one connectivity. (c) Brain mapping of the FC variations due to the participant, within-subject, residual, and scanner factors. (d) Brain networks of the FC variations summarized using Yeo’s 7-network parcellation. All reported values are represented by Fisher’s z-transformed Pearson correlation coefficients.

Subsequently, we examined brain regions and networks affected by each factor. A trend similar to that of the BMB dataset was observed, with some exceptions (Fig. 3(c), (d)). The within-subject variation was large in terms of connectivity involving the cerebellum and visual cortex, and large scanner-related variation was observed in the orbito-frontal cortex. The pair-wise scanner distance matrix and dendrogram computed on the basis of the agglomerative hierarchical clustering revealed that the Phillips Achieva and Siemens scanners were similar; however, a larger separation was observed for the Signa and MR750W scanners from GE and Siemens scanners (Supplementary Fig. 2(c)).

### FC differences between neuropsychiatric disorder groups and heathy controls (HCs)

We examined group-level FC differences between patients with MDD, ASD, or SCZ, and HCs to compare the FC variations from the disorder-unrelated factors, e.g., imaging protocol, scanner, participant, and within-subject variations.

The magnitude distributions of the FC differences for each disorder are shown in Fig. 4(a). The median values (5^th^–95^th^ percentile) of the magnitudes for the MDD, ASD, and SCZ groups were 0.019 (0.002–0.061), 0.020 (0.002–0.062), and 0.029 (0.003–0.086), respectively, which were as small as the FC variations of the scanner and imaging protocol factors. However, examining the upper tails of each distribution revealed substantial effects for certain connections; more specifically, approximately 0.5%, 0.3%, and 2.3% of all connections exhibited a magnitude exceeding 0.1 in the MDD, ASD, and SCZ groups, respectively, highlighting the fact that accurate biomarker development requires the selection of important disorder-specific connections. Furthermore, the comparatively smaller FC differences in the MDD and ASD groups in comparison with that of the SCZ group were indicative of the challenges associated with creating precise biomarkers for MDD and ASD. The comparison of the disorder-related and disorder-unrelated FC variations from the BMB traveling-subject dataset for the 50 largest disorder-related connections (Supplementary Figs. 3–5) revealed that the magnitudes of the scanner and imaging protocol factors were small for most connections, whereas those of the within-subject and participant factors were as large as the disorder-related differences.

**Figure 4.**
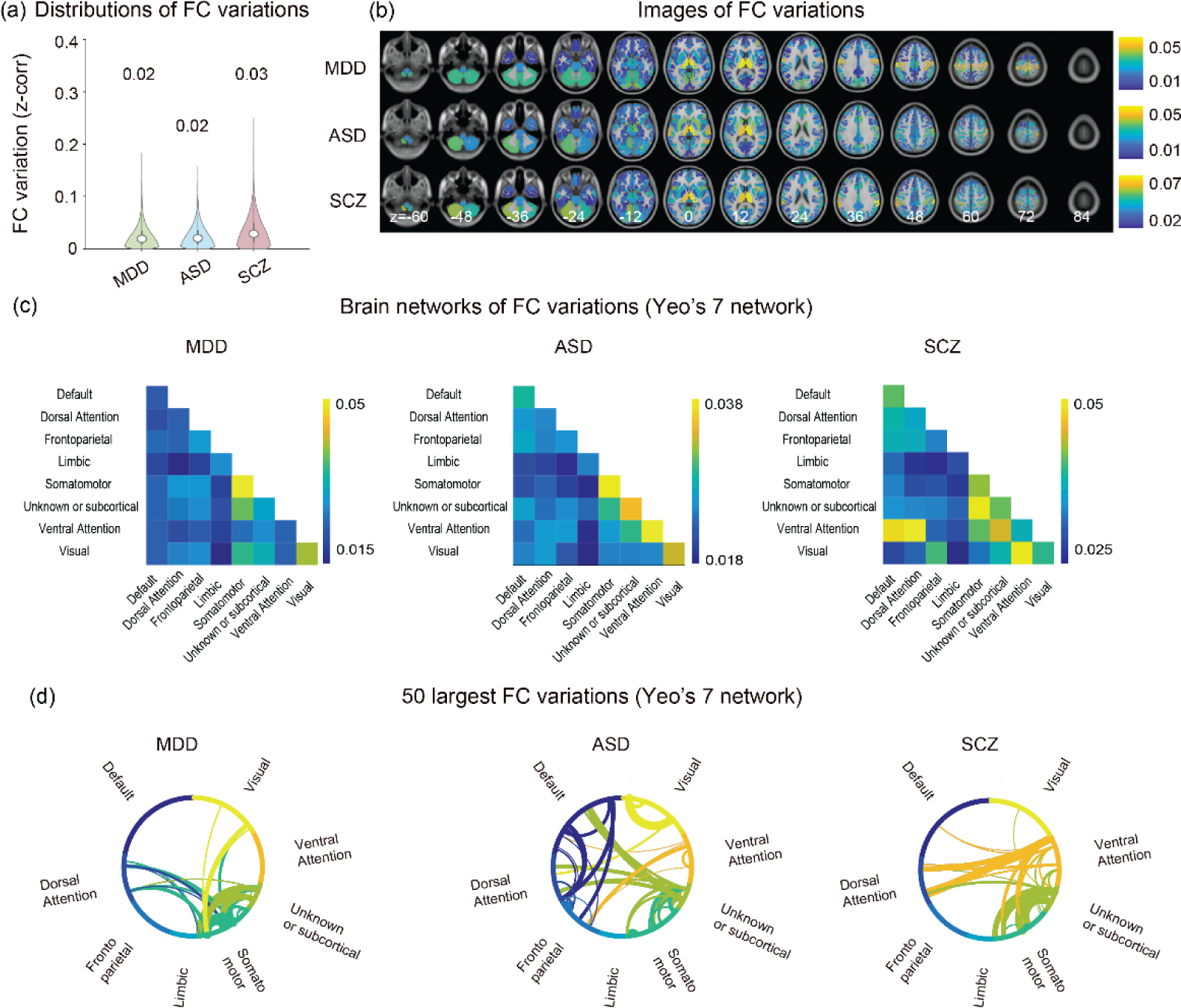
Comparison of the disorder-related functional connectivity (FC) differences between groups of patients with major depressive disorder (MDD), autism spectrum disorder (ASD), or schizophrenia (SCZ) and age- and-sex-matched heathy controls. (a) Distributions of the magnitudes of the disorder-related FC differences. (b) Brain mapping of the disorder-related FC differences (c). Brain networks of the FC differences summarized using Yeo’s 7-network parcellation. (d) The 50 largest FC variations summarized using Yeo’s 7-network parcellation. All reported values represent Fisher’s z-transformed Pearson correlation coefficients.

The brain regions and networks affected by each disorder are presented in Fig. 4(b), (c). For all three, a large magnitude was observed for connectivity involving the thalamus. For the MDD group, in addition to the thalamus, large FC differences were observed in the somatosensory and motor regions, consistent with the results of a recent extensive data analysis conducted by the PsyMRI consortium^32^; however, this contradicts early studies that emphasized default mode and front-parietal network involvement^30,39^. For the ASD group, large FC differences were observed for intra-network connections involving somato-motor, ventral attention, subcortical, and visual networks. For the SCZ group, large FC differences were observed in the thalamus and somatosensory and motor regions in addition to changes in inter-network connections involving the ventral attention network.

Collectively, these results indicated that the impact of the scanner and imaging protocol factors on biomarker development could be limited, whereas within-subject and participant factors could have a greater impact and may require more careful consideration.

### Signal-to-noise ratio (SNR) enhancement using multivariate connectivity biomarkers

We have shown that individual differences (participant factor) and within-subject variations were as large as or even larger than the disorder-related differences at most connections, highlighting the difficulty of developing reliable univariate connectivity biomarkers that permit individual-level classification. However, previous studies involving machine-learning algorithms have demonstrated that *multivariate* connectivity biomarkers can facilitate individual-level classifications^17,19,40^. To understand the factors affecting such outputs, we investigated the impact of variations caused by disorder-unrelated factors in the BMB traveling-subject data on the MDD, ASD, and SCZ biomarkers we developed in a previous study using ensembles of least absolute shrinkage and selection operator (LASSO) classifiers^19^. We hypothesized that machine-learning algorithms could optimize connectivity weightings to empower the model to distinguish patients from HCs, while suppressing the influence of individual differences within the patients and HCs group.

First, we investigated the connections selected by the LASSO algorithm using the magnitude distributions of the FC variations. For comparison, we consider the naïve greedy strategy in which the 50 largest disorder-related connections (see Supplementary Figs. 3–5 for details of each individual FC) were selected and averaged. For the MDD biomarker (Fig. 5(a)), the FC variation distributions of the top 50 LASSO-selected connections (histogram left-half) were significantly different from those of the 50 largest disorder-related connections (histogram right-half) with respect to the participant, within-subject, scanner and disorder factors (two-sample Kolmogorov-Smirnov test, p<0.001). The LASSO algorithm did not limit its selection to the functional connections with the 50 largest differences between the disorder and HC groups; instead, it selected those with smaller individual-, within-subject-, and scanner-related variations. Compared with the magnitude distributions of all 71,631 FCs (violin plots), the distributions of the lasso-selected FCs had the significantly different distribution only for the disorder factor (two-sample Kolmogorov-Smirnov test, p<0.001). On the other hand, the distributions of the greedily selected FCs had the significant difference for the participant and within-subject factors as well as the disorder factor (two-sample Kolmogorov-Smirnov test, p<0.001). Similar results were observed for the ASD and SCZ biomarkers, except for the lack of significant differences in scanner-related FC variations (Fig. 5(b)(c)).

**Figure 5.**
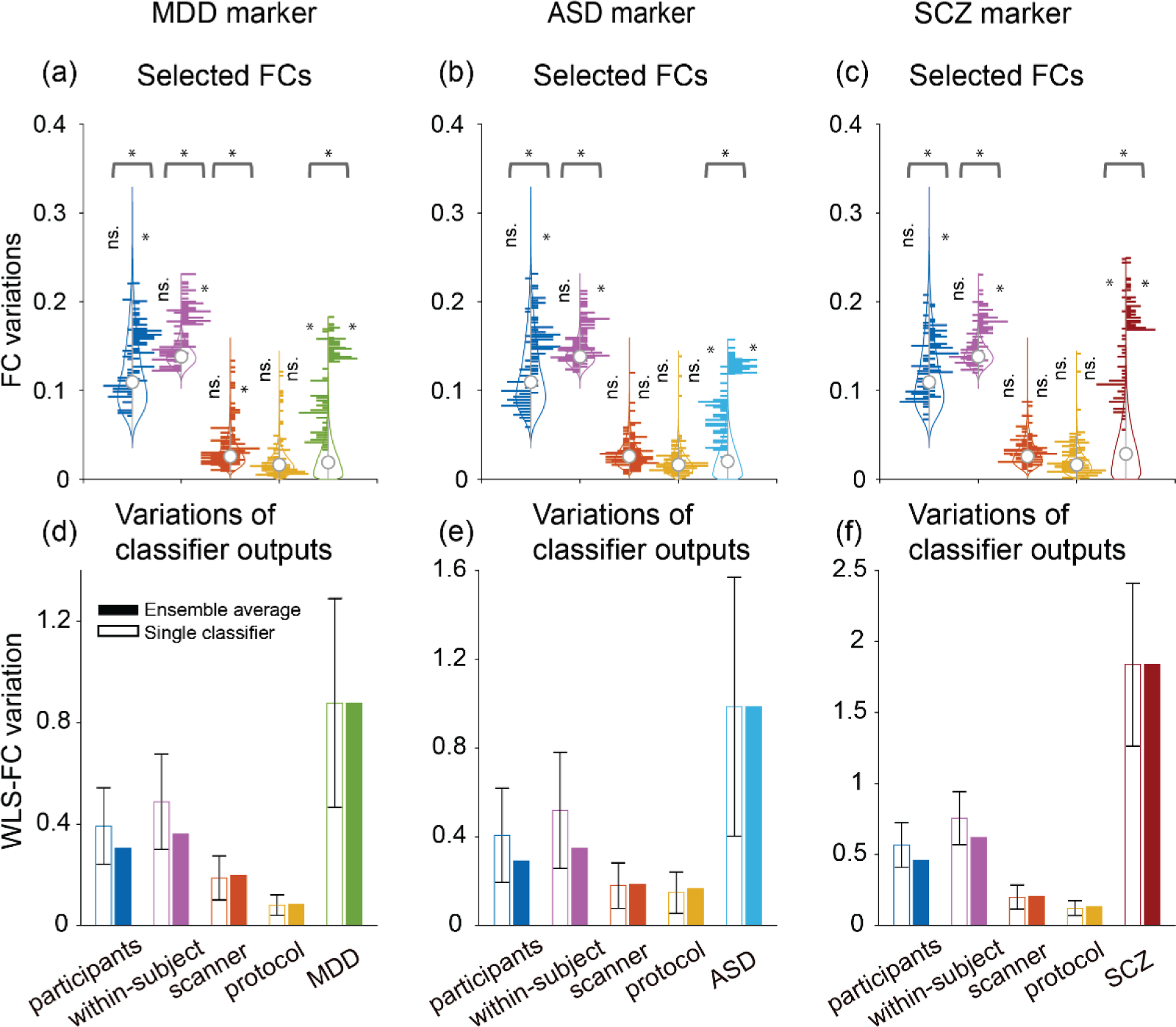
Analysis of the effects of functional connectivity (FC) variations on the multivariate connectivity biomarker outputs. We evaluated the impact of FC variations due to the participant, imaging protocol, scanner, within-subject, and disorder factors on the outputs of major depressive disorder (MDD), autism spectrum disorder (ASD), and schizophrenia (SCZ) biomarkers developed in our previous study^19^. The biomarker for each psychiatric disorder consisted of an ensemble of 100 linear classifiers, each of which was trained with partially overlapping but distinct subsampled data using the least absolute shrinkage and selection operator (LASSO) algorithm. The output of each classifier was a scalar value that represented the weighted linear summation of the FC (WLS-FC). The final decision value, indicating the likelihood of the presence of the disorder, was obtained by averaging the outputs of all 100 classifiers. (a) The distributions of the magnitudes of the FC variations of 50 LASSO-selected connections (left-half histogram) and the top 50 MDD-related connections (right half histogram, for the connections displayed in Fig. 4(d)) superimposed on the magnitude distributions of all 71,631 FCs (violin plots, integrated with Fig 2(a) and 4(a)). The asterisk and ns. indicate statistical significance and non-significance (p < 0.001) from two-sample Kolmogorov-Smirnov test, respectively. (b)(c) The same data analyses are shown for the ASD and SCZ biomarkers. (d) The impact of the FC variations on the output of the MDD biomarker. The unfilled bars represent the variation of each classifier output averaged across all 100 classifiers (the error bar indicates the standard deviation), whereas the filled bars represent the variation of the final decision value. (e)(f) The same analyses are shown for the ASD and SCZ biomarkers, respectively.

Subsequently, we evaluated the weighted linear summation-FC (WLS-FC) variations caused by the disorder-unrelated and disorder-related factors to assess their impact on the decision value of the diagnostic biomarkers. The WLS-FC variations for the MDD biomarker are presented in Fig. 5(d); the mean variations of each individual classifier output (shown by the unfilled bars) caused by the participant and within-subject factors were less than half of the difference between the MDD and HC groups, whereas variations caused by scanner and protocol factors were much smaller (approximately one-tenth of that difference). Further reduction in WLS-FC variations was observed for the ensemble-averaged output (filled bars), particularly for the participant and within-subject factors. The same tendency was observed for the ASD and SCZ biomarkers (Fig. 5(e), (f), respectively).

To quantify the signal improvement, we estimated the SNR of each FC or the classifier output as the disorder-related difference divided by summation of the participants, within-subject, scanner and imaging protocol variations (Fig. 6 and Supplementary Table 1). For MDD, the distribution of the SNR estimates of each individual FC from all 71631 FCs ranged from 3.8 × 10^−7^ to 0.48 with the median value 0.064. The average SNR of top 50 disorder-related FCs and top 50 LASSO-selected FCs were 0.365 (±0.063) and 0.256 (±0.095), respectively. The SNR estimates of the LASSO classifier outputs before and after ensemble average were 0.742 (±0.132) and 0.965, respectively. Thus, the ensemble LASSO classifier improved the SNR by 15 times and 2.6 times compared with the median SNR of whole 71631 FCs and the greedy univariate strategy. The similar values of the signal improvement were observed for ASD and SCZ (ASD: 15.4 times and 3.1 times, SCZ 13.9 times and 2.6 times). In summary, improvements of SNRs by the ensemble LASSO biomarkers were achieved by the following three mechanisms: first, selecting FCs with large disorder effects and modest participant or within-subject variability, second, spatially weighted averaging of the selected FCs reduced the participants and within-subject variations significantly, and third, the ensemble of 100 LASSO classifiers further reduced the variations. We here roughly quantify contributions by the three factors in improving the SNRs for the MDD. First, the top 50 LASSO-selected FCs have 0.256/0.064=3.9 times larger than the median SNR of whole 71,631 FCs. Second, the linear weighted summation of the LASSO classifier improved 0.742/0.256=2.9 times by weighted spatial averaging; this is reminiscent of replacing spatial averaging by temporal averaging based on Ergodic property in statistical physics, but the opposite in our case, that is, replacing temporal averaging by spatial averaging. Third the ensemble averaging improved 0.965/0.742=1.3 times. Altogether, the entire procedure improved SNR by 15 times. This is in sharp contrast to SNRs close to but less than 1 for the 50 largest disorder-related FCs. The ASD case is similar to MDD, and the SCZ case attains a little higher SNR because of relatively larger disorder effects.

**Figure 6.**
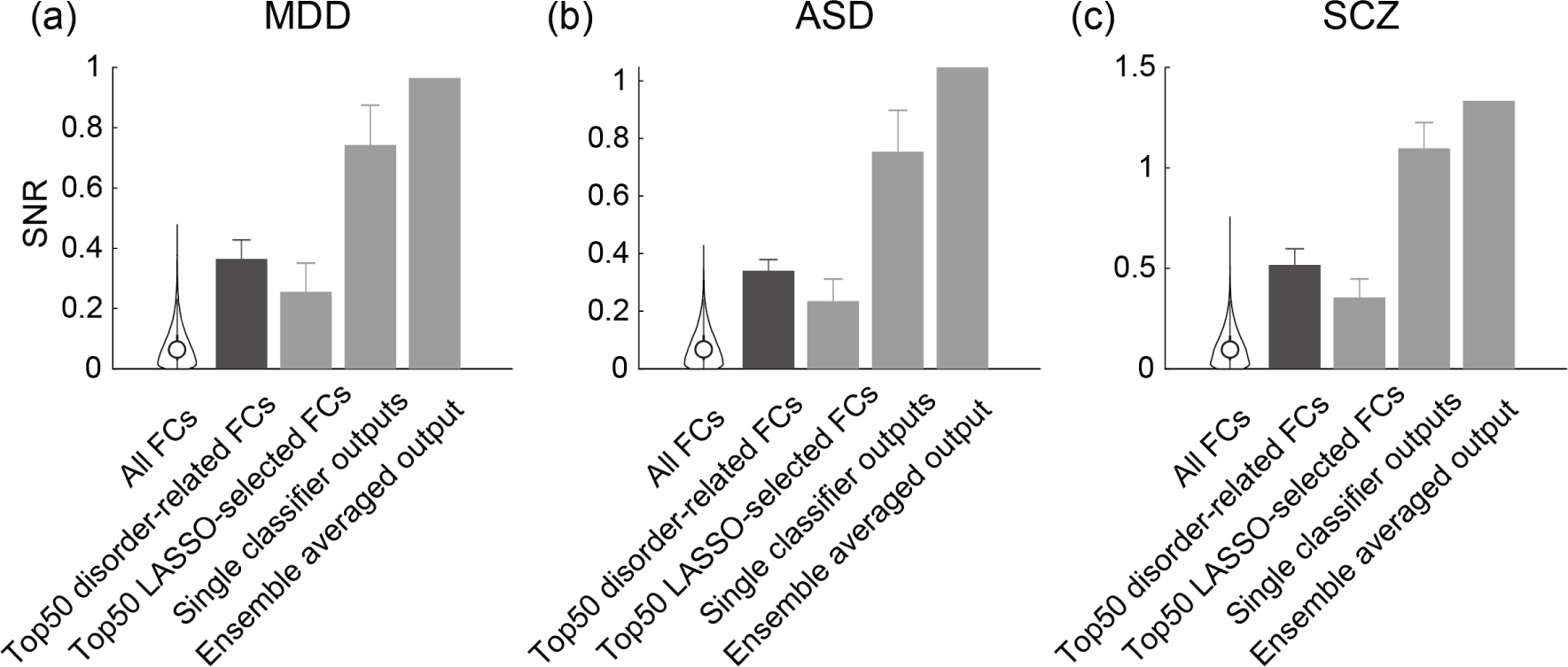
Signal-to-noise ratio (SNR) estimates of the FCs and the LASSO classifier outputs for MDD, ASD and SCZ. For each individual FC or classifier output, the SNR was defined as the disorder-related variation divided by summation of the participants, within-subject, scanner and imaging protocol variations. ‘All FCs’ represents the SNR distribution of the individual FCs collecting from all 71631 FCs. ‘Top 50 disorder-related FCs’ represents average SNR of the 50 individual FCs with the largest MDD-related variation. ‘Top 50 LASSO-selected FCs’ represents average SNR of the 50 individual FCs which was most frequently selected by the LASSO classifiers. ‘Single classifier outputs’ and ‘Ensemble averaged output’ represent the SNRs of the LASSO classifier outputs before and after ensemble average, respectively. (a)(b)(c) the SNR estimates of MDD, ASD and SCZ, respectively. The SNR values are listed in the supplementary table 1.

These results supported our hypothesis that machine-learning algorithms could optimize the connectivity weighting to maximize disorder-related differences while simultaneously suppressing individual differences. Contrary to our expectations, we observed suppressed within-subject variations as well.

## Discussion

The present study involved a comprehensive and quantitative evaluation of the effects of various factors on FC and machine learning-based biomarker outputs to delineate directions for future research to establish reliable individual-level biomarkers. The carefully-designed linear fixed modeling of the traveling-subject datasets revealed that the effects of individual differences and residual components on FC variations were several times greater than the variations caused by scanner and imaging protocol factors. Additionally, the FC variations attributed to the residual components were highly similar to the within-subject across-runs variations. Each factor affected brain regions and networks differently; for example, large individual FC differences were observed in the dorsal attention and front-parietal networks; large within-subject FC variations were observed in the sensory and motor-related regions; and large scanner- and imaging protocol-related FC variations were evident at the bottom and top of the brain. The disorder-related FC difference was as small as the scanner- and imaging protocol-related variation on average and was of similar magnitude to the individual FC differences and within-subject FC variation at the connectivity with the largest disorder effects. By evaluating the variations of the multi-connectivity biomarker outputs, we found a reduction of the individual difference and within-subject variations by optimal weighting of multiple connectivity and ensemble averaging. Our results revealed large variability in disorder-unrelated factors at the level of single connections; however, the use of multivariate connectivity biomarkers served as a noise-suppression mechanism, increasing the SNR through optimal weighting and ensemble averaging. Furthermore, this study was the first to evaluate FC variations related to specific imaging protocols and their effects on biomarker outputs. All of our reported values for the FC variations were based on Fisher’s z-transformed Pearson correlation, enhancing interpretability across studies. This approach differs from previous traveling-subject studies, offering a more comprehensive understanding of FC variations and their implications on imaging biomarker development ^35,36^.

The linear fixed effects modeling revealed that the FC variation caused by the participant factor was several times larger than that caused by scanner and imaging protocol factors, with even larger variation for the unexplained residual components. The distribution for the participant variation was broad across connections, whereas the distributions for the other three types of variations were narrower across both traveling-subject datasets, with similar outcomes observed when the FC values were derived from volumed-based anatomical parcellation (BrainVISA atlas, 137 regions without the cerebellum; Supplementary Figs. 7 and 8). The larger contributions of the participant factor relative to that of the other measurement factors were also consistent with findings of two previous traveling-subject studies^35,36^. This study was also the first to incorporate the decomposition of measurement bias into the imaging protocol and scanner factors, revealing slightly larger scanner-related FC variation than imaging protocol-related FC variation. The larger impact of the participant factor in the BMB dataset compared with that of the SRPBS dataset could be due to the wider variability in subject demographics in the former dataset. For example, the SRPBS traveling-subject dataset included only young, adult males, whereas the BMB dataset included both male and female adults in their twenties to sixties.

The largest FC variations were associated with residual components that could not be explained by the linear fixed effects model. The ratios of the unexplained residual variations to the total variations averaged across all connections were 69% and 64% for the SRPBS and BMB datasets, respectively, consistent with the 60–80% ratios attributed to unexplained residual components in a previous study^35^. However, the previous traveling-subject studies did not investigate the residual components in detail. The present study clearly showed the relevance of the residual components with the within-subject variations by high connectivity pattern similarity between them (Fig. 2,3(a)). This result remained unchanged when the data used for computing the residual components and within-subjects variation were completely separated (Supplementary Fig. 9(a)). These results suggested that the large proportions observed for the unexplained residual components reflected the within-subject FC variation. It is important to note that the within-subject variations here include variations across different days, as each participant’s data comprised at least two days of experiments, on average approximately 8 weeks apart, with three runs per day. The within-day variation for a participant can be determined by subtracting the day-specific FC pattern, averaged over three runs, from each run’s data. We discovered that the within-participant within-day FC variation shared a similar connectivity pattern with the current study’s intra-participant inter-run FC variation, albeit with a reduced magnitude (Supplementary Fig. 10).

The brain mapping of the participant, imaging protocol, scanner, and within-subject FC variations revealed distinct patterns in the brain regions affected by each factor, with some overlap. For example, the participant FC variations were large in regions associated with the dorsal attention and frontoparietal networks, whereas the within-subject FC variation was large in the somatosensory, motor, and visual cortices, as well as in certain regions within the dorsal attention network. These distinct patterns were observed in both the BMB and SRPBS datasets, with an exception being the large within-subject FC variation observed in the cerebellum and visual cortex only in the SRPBS dataset. Similar distinct patterns of between-subjects and within-subject FC variations have been reported in several previous studies^41,42^. Large imaging protocol- and scanner-related FC variations were predominantly observed on the top and bottom of the brain, although the detailed patterns slightly differed for each factor. A large imaging protocol FC variation was observed in the anterior and inferior parts of the brain, including the orbitofrontal cortex, gyrus rectus, and olfactory regions, whereas large scanner-related FC variation was observed in the superior frontal gyrus and cerebellum; the latter FC variations were also high in the anterior frontal parts of the brain for the SRPBS dataset, resembling the scanner differences reported previously^35^.

The comparison of the FC variations between the disorder-related factor and the participant, within-subject, scanner, and imaging protocol factors (Fig. 4 and Supplementary Figs. 3–5, 11) has particularly important implications for psychiatric biomarker development. First, a small subset of connections exhibited substantial group differences between patients and HCs. For example, the number and magnitude of functional connections relevant to MDD and ASD were comparatively smaller than those relevant to SCZ, suggesting that developing accurate MDD and ASD biomarkers might be more challenging. Second, the disorder-related FC differences were comparable to or smaller than the individual difference and within-subject variations, even when focusing on 50 connections with the largest disorder-related differences (Supplementary Figs. 3– 5). Aggregating multiple connectivity changes is essential to allow for differentiating patients from HCs at an individual level. Third, the magnitude of the imaging protocol-related FC variations was small for most of the MDD-, ASD-, and SCZ-related connections, except for several MDD-related connections around the somatosensory and motor regions. Thus, imaging protocol-related FC differences may have a limited impact on biomarker development. This finding is particularly important because it suggests possibility of integrating two data sets from two distinct nationwide projects SRPBS and BMB, enabling machine-learning-based biomarker development using combined datasets comprising approximately 10,000 samples.

The investigation of the effect of disorder-unrelated variations on the multi-connectivity biomarker outputs that we previously developed using the ensemble LASSO algorithm revealed that individual difference and within-subject FC variations could be reduced by optimal weighting of automatically selected FCs and ensemble averaging. The reduced effect attributed to individual differences was expected, as the machine-learning algorithm attempts to reduce within-group variance while simultaneously increasing the between-group variance (i.e., differences between disorder and HC groups). However, the reduced impact of the within-subject variation was surprising because there was no explicit source of information about the such variation that the machine-learning algorithm could have used in the training data, that is, no multiple scans from individual participants were included in the discovery cohort for the biomarker development; however, this effect might have been because the within-subject variations had some commonality across subjects. Even though the individual difference and within-subject variations were reduced, the magnitude of these two factors was larger than that of the scanner and imaging protocol variations. This observation underscores the challenge of further reducing the influence of individual differences and within-participant variations to create even more robust and precise biomarkers, which is especially critical for precision medicine applications. Addressing this challenge will require innovative experimental and analytical approaches in future research endeavors. One possible progression involves integrating the SRPBS and BMB datasets to form big datasets for biomarker discovery. The expanded dataset size would allow for an increase in the number of the selected FCs by the ensemble sparse classifiers. This increased number of FCs would boost the SNR in distinguishing disease effects from irrelevant variations, resulting in biomarkers that are more reliable, generalizable, precise, and applicable.

This study had several limitations. First, the use of the statistical model for factor decomposition is limited to linear modeling with only a few factors. Although it may be desirable to include non-linear effects, such as interactions between imaging protocols and scanner types or to include other factors, such inclusion complicates the explanatory matrix, resulting in the null space being impossible to interpret. Despite our efforts to account for data variance with known factors, substantial portions remained unexplained. Second, the SNR values in Fig. 6 and Supplementary Table 1 were only rough estimates. We assumed the four disorder-unrelated factors as noise as well as independence among the four factors. In addition, our estimates of signal could be overestimated because the same dataset was used for both developing the classifiers and evaluating disorder-related differences of the biomarker outputs. However, this overestimation is not expected to affect the SNR comparison between the LASSO biomarkers and the top 50 disorder-related FCs. Third, our analysis was based on 10-minute rsFC trials. The experiment duration significantly affects the test-retest reliability of rsFC^42,43^, so the extent of within-participant variation should decrease with longer trial durations. Forth, we did not implement a preprocessing pipeline specifically optimized for the HARP and CRHD protocols. More sophisticated preprocessing techniques^38,44^, which have been proposed for data acquired with these protocols, could further improve biomarker performance. The impact of preprocessing on FC variations should be explored in future studies.

In conclusion, this study provided comprehensive and quantitative understanding of the effects of various factors on FC and machine-learning–based biomarker outputs. Our study also demonstrates the benefit of characterizing each FC variation from diverse perspectives, encompassing not only disorder-related differences but also disorder-unrelated variations, e.g., those attributed to participant, within-subject, imaging protocol, and scanner factors. This comprehensive approach is instrumental in advancing the development of more robust, generalizable, and accurate biomarkers.

## Methods

### Datasets

This study analyzed two traveling-subject datasets derived from two national-wide projects conducted in Japan (SRPBS, 2012–2018 and BMB, 2018–2023), as well as a portion of the SRPBS multi-disorder dataset. The SRPBS project was the pioneering multicenter study conducted to develop multicenter generalizable psychiatric biomarkers using a unified imaging protocol. The subsequent BMB project aimed to improve biomarker selection using a cutting-edge imaging protocol and data processing techniques. To bridge the biomarker selection between two projects, the BMB traveling-subject data were acquired using the SRPBS imaging protocol and two new imaging protocols. Investigating the effects of these imaging protocols on FC variability was one of the focuses of the present study.

A comprehensive description of the SRPBS traveling-subject and multi-disorder datasets was provided in our previous study^37^. All of SRPBS data used for the analysis is publicly available. To summarize, nine young adult male subjects (age range: 24–32 years; mean age: 27±2.6 years) visited 12 sites in the traveling-subject dataset, participating in fMRI experiments involving two or three runs of 10-minute resting-state testing within a single experimental session at each site, resulting in the acquisition of 411 runs of 10-minute eyes-open resting-state fMRI data. We employed a unified imaging protocol referred to as the SRPBS protocol. Owing to hardware limitations, two phase-encoding directions were used depending on the scanners used, either A->P (anterior to posterior) or P->A (posterior to anterior). However, we did not differentiate the phase-encoding direction as different imaging protocols because we corrected for the impact using the corresponding field maps. The dataset included data collected from seven types of scanners from three different MRI manufacturers (Siemens, GE, and Phillips). The SRPBS multi-disorder dataset used for the analysis consisted of data from approximately 900 subjects, acquired using the SRPBS imaging protocol at the following four sites. The dataset included data from patients with three psychiatric disorders (MDD, ASD, and SCZ) and HCs (see Supplementary Figs. 12 and 13 for graphical representations of the data).

Specific details pertaining to the BMB dataset were described in our previous study^38^. Briefly the data were derived from 75 subjects (48 males and 27 females; mean age: 31.8±10.0 years) from 17 sites. Each subject had visited three or more sites, including one of three hub sites according to a hub-and-spoke model, which differed from the SRPBS traveling-subject design in which all subjects visited all sites. For each participant, data were collected from at least two runs of a 10-minute eyes-open resting-state fMRI task conducted in a single experimental session at each site. At least five healthy participants were recruited at each site. In total, approximately 1,200 runs of 10-minute eyes-open resting-state fMRI data were obtained using three imaging protocols, including the SRPBS protocol previously mentioned, the CRHD protocol, which is the MRI protocol developed by the CRHD initiative of the Human Connectome Project (HCP) in the United States of America (USA), customized for high-performance, 3T MRI scanners such as the MAGNETOM Prisma (Siemens Healthcare GmbH, Erlangen, Germany), and the HARP, which is an HCP-style protocol with a short scanning time optimized for clinical studies so that it can be used for multiple MRI scanners/sites and was designed to obtain high-quality, standardized brain MRI data in a ‘clinically’ practical time window (see Supplementary Table 2 for more details). Seven scanner types from two MRI manufacturers (Siemens and GE) were included (see Supplementary Fig. 14 for a graphical representation of the data).

All participants in all datasets provided written informed consent, and all recruitment procedures and experimental protocols were approved by the institutional review boards of the principal investigators’ respective institutions.

### FC computation

We computed a region-level whole-brain FC matrix using identical processing steps for both the SRPBS and BMB datasets.

The resting-state fMRI images underwent preprocessing using the standard pipeline implemented in fMRIPrep 1.0.8,^45^ which consisted of several steps, including the exclusion of the initial 10 seconds of data for T1 equilibration, slice-timing correction, realignment, coregistration, distortion correction using a field map, segmentation of T1-weighted structural images, normalization to the Montreal Neurological Institute space, and surface projection.

Subsequently, the resting-state fMRI timeseries data underwent the following processing steps: physiological noise removal by employing 12 regressors, which consisted of six head motion parameters, signal averaging across the entire brain, and five anatomical component-based noise correction (CompCor) components. The data were then bandpass-filtered using a second-order Butterworth filter with a passband ranging from 0.01 to 0.08 Hz. Additionally, image volumes affected by head motion, as indicated by a frame displacement exceeding 0.5 mm^46^, were eliminated from further analysis, as were runs with excessive head motion. A region-level whole-brain connectivity matrix was computed using Glasser’s surface-based MMP atlas; this consisted of 379 regions of interest (ROIs) (360 cortical parcels and 19 subcortical parcels)^44^ using the ciftify toolbox version 2.0.2–2.0.3. The region timeseries was obtained by averaging the voxel timeseries within each region. The connectivity matrix was obtained by calculating the Pearson correlations between all regional timeseries pairs. Since the connectivity matrix is symmetric, the lower triangular elements were extracted, and a vector was formed with a size of 71,631×1 (referred to as a connectivity vector). Finally, Fisher’s z-transformation was applied to each element of the connectivity vector.

After excluding any data affected by image processing errors, excessive head motion, and an insufficient number of runs at a single site from further analysis, we analyzed the connectivity vectors from a total of 398 runs from 12 sites, including nine subjects, one protocol and seven scanner types in the SRPBS traveling-subject dataset, and the connectivity vectors from a total of 1,167 runs from 14 sites, including 73 subjects, three protocols, and six scanner types in the BMB traveling-subject dataset (see Supplementary Tables 3 and 4 for more details).

### Estimation of FC variations due to experimental factors

To determine the influence of experimental factors such as the participant, scanner, or imaging protocol on FC, we used a linear fixed effects model for each connection, which allowed us to estimate the magnitude of these factors’ effects. We used a three-factor model consisting of the participant, scanner, and imaging protocol for the BMB traveling-subject data, whereas a two-factor model consisting of the participant and scanner factors was used for the SRPBS traveling-subject data.

We let *z*_*nc*_ be a z-transformed connectivity strength for a specific run indexed by n and a connection indexed by c (N: total number of runs, C: total number of connections) and let 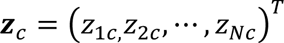 be a column vector containing all the strength for connection c across all runs. Then, in the case of the three-factor model, we assumed a linear regression model with three explanatory variables was represented as,

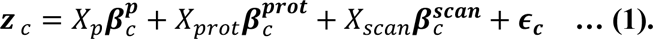

Here, three factors are categorical values represented by the binary matrices *X_p_*, *X_prot_* and *X_scan_*, respectively. For example, the participant-factor matrix *X_p_* would be a matrix of size N-by-P (with P being the total number of participants) and the *ij*th element would be 1 if the participant indexed by *i* participated in a run indexed by *j*; otherwise, it would be 0. The parameter vector 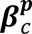 was a vector of size P-by-1 whose element represents the magnitude for each participant. The explanatory matrices and the parameter vectors for the protocol and scanner factors were defined in the same way. The term *ϵ_C_* represented residuals that cannot be explained by the linear summation of three factors.

The equation (1) can be rewritten in the simplified form

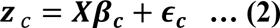

where 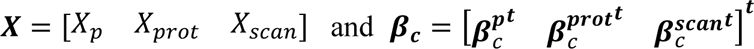 are a matrix and vector concatenating three factors, respectively. Using the least squares method, the parameter vector ***β****_c_* could be obtained by solving the following normal equation,

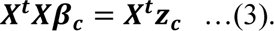

Since all three factors were categorical variables, the concatenated explanatory matrix ***X*** was not of full rank, which could be easily confirmed by summing the columns of *X_p_*, *X_prot_* and *X_scan_*, where all results would be a vector consisting of all ones. Thus, the linear equation (3) did not have a unique solution (and the inverse of ***X^t^X*** does not exist), but the least squares solution could be obtained using the Moore-Penrose generalized inverse matrix,^47^

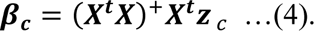

This was the solution of the linear system (3) in which the minimum L2-norm and the null space of ***X^t^X*** could be determined from the design of the explanatory matrix. The singular value decomposition of the explanatory matrix provided information about the null space. Thus, in our three-factor or two-factor model, we confirmed that the undetermined components were constant values within each factor (see Supplementary Fig. 9 for more details). Thus, the baseline values of each factor were arbitrary, and only the differential values within each factor were meaningful.

### Quantification of FC variations due to experimental factors

We computed the FC variations attributed to the participant (or individual subject), protocol, and scanner factors by determining the standard deviation of 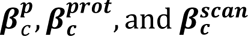 across the members within each factor, respectively. The residual FC variation was obtained as the standard deviation of *ϵ_C_*. The pair-wise distance matrix between members of the scanner type and imaging protocol factors was computed by calculating the mean absolute difference between corresponding estimated parameters across all connections.

### Computation of within-subject across-runs FC variations

To investigate the origin of residual FC variations resulting from the linear fixed effects modeling, we computed within-subject across-runs FC variations directly from the FC vectors. The within-subject FC variation is typically defined by the variability of connectivity patterns between different runs within one subject. In this study, the within-subject FC variations averaged over screened subjects were defined as follows. First, we screened for subjects who had performed at least six runs under a single measurement condition, such as the same site, protocol, or scanner, to estimate the subject-specific within-subject FC variations robustly. For each chosen subject, the data were collected from at least two days of experimental sessions, with three runs per day, and the subject-specific within-subject FC variations were obtained by determining the connectivity-wise standard deviation of the connectivity vectors across runs for that particular subject. Then, the subject-average within-subject FC variations, which we simply referred to as the within-subject FC variations in this study, were computed as the average of all subject-specific within-subject FC variations.

After screening subjects with at least six runs under a single measurement condition, the SRPBS dataset contained 132 runs from nine subjects acquired using a Trio scanner at a single site named ATR, whereas the BMB dataset contained 201 runs from 31 subjects acquired with the HARP and four different scanner types across seven sites.

### Computation of disorder-related FC variations

For clinical applications, it is crucial to compare the FC differences associated with neuropsychiatric disorders with those that are unrelated to the disorder, such as those associated with the imaging protocol or scanner, or within-subject and participant FC variations identified in the traveling-subject data analysis. Therefore, we computed the disorder-related FC differences of three psychiatric disorders, including MDD, ASD, and SCZ, using a portion of the data from the SRPBS multi-disorder dataset. The whole-brain FC matrices were computed using Glasser’s MMP atlas in the exact same way as for the traveling-subject datasets. The statistical harmonization using the SRPBS traveling-subject data was applied to reduce the site effects^30^. For each psychiatric disorder, we randomly selected HC subjects from the dataset for age-, sex-, and site-matching as much as possible, with 138 patients with MDD and 138 HCs (age: 42.12 ± 12.33 *and* 41.76 ± 12.39 *years*, *respectively*; male ratios: 0.46, 0.54); 109 patients with ASD and 109 HCs (age: 29.14 ± 8.35 *and* 31.25 ± 7.33 *years*, *respectiely*; male ratios: 0.84, 0.87), and 84 patients with SCZ and 84 HCs (age: 37.20 ± 11.24 *and* 37.18 ± 11.46 *years*, *respectively*; male ratios: 0.58, 0.57). The disorder-related FC differences were computed as the absolute value of the group-averaged FC difference between the patient and matched HC groups for each disorder.

### Analysis of the FC variations on multivariate connectivity biomarker outputs

To understand the underlying mechanism behind the effectiveness of the multivariate connectivity biomarkers on individual-level classifications, we analyzed the variations attributed to the disorder-unrelated and disorder-related factors on the biomarker outcomes. More specifically, we evaluated the variations attributed to the participant, imaging protocol, scanner, and within-subject variation from the BMB traveling-subject dataset and disorder factors from the SRPBS multi-disorder dataset using the weight parameters of MDD, ASD, and SCZ biomarkers we identified previously ^19^. The biomarker of each psychiatric disorder consisted of an ensemble of 100 linear classifiers, each of which was trained using partially overlapping but distinct subsampled data using the least absolute shrinkage and selection operator (LASSO) algorithm and FC vectors computed with the MMP atlas. The output of each classifier was a scalar value that indicated the likelihood of the presence of the disorder, with the final decision value obtained by averaging the outputs of all 100 classifiers. To clarify, if we denote the weight parameter of the n-th classifier by **W_n_** and the FC vector of a subject by **X**, then the final decision value was calculated as 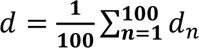, where 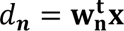 represents the decision value of each individual classifier. Thus, each classifier output is a WLS of multiple FC values, which was referred to as the WLS-FC.

First, we investigated the FCs selected by the LASSO algorithm using the magnitude distributions of the FC variations to assess the feature selection preferences of the machine-learning algorithm. Given that the LASSO algorithm performs the feature selection during the optimization of the parameter weighting, we focused on the frequently selected FCs among the 100 classifiers and chose the top 50 most frequently selected FCs on the basis of the average numbers of FCs selected for the MDD, ASD, and SCZ biomarkers (49, 54.2, and 53.7, respectively). For comparison, we conducted two-sample Kolmogorov-Smirnov testing between the FC variation distributions consisting of the LASSO-selected connections and those consisting of the 50 largest disorder-related connections (Fig. 5(d)).

Second, we analyzed the WLS-FC variations (i.e., variations of the biomarker outcomes) attributed to the participant, scanner, imaging protocol, within-subject, and psychiatric disorder factors, which were calculated for the output of each individual classifier and the output after ensemble averaging. To compute the WLS-FC variation associated with a particular factor, we required the FC deviation vector for each member of the factor. Each element of the FC deviation vector was defined by a signed scalar value representing the deviation from the mean across the members of the factor. For example, the FC deviations of the participant factor were computed by subtracting participant-averaged beta estimates from each participant for each FC. The imaging protocol- and scanner-related FC deviations were computed in the same way. The disorder-related FC deviation was computed as the difference between the group averages. The within-subject FC deviation was computed by pooling the within-subject FC deviations of each subject, each of which was obtained by subtracting the run-averaged FC from each run’s FC data within a subject. Subsequently, we calculated the WLS-FC variations by taking the standard deviation of the WLS-FC deviations, which was obtained by projecting FC deviation vectors on the biomarker space defined by the classifier weight parameters.

## Acknowledgments

This study was supported by AMED (grant numbers: JP19dm0207069, JP18dm0307001, JP18dm037002, JP18dm0307004, JP23dm0307008, JP23dm0307009, JP21dm0207070h0003, JP21dm0307003h0004, and JP21dm0307004h0004), JST MS9 (grant number: JPMJMS2291), and JSPS KAKENHI (grant number: JP19H05726 and 23H00414).

## Author Contributions

Conceptualization: OY and MK

Methodology: OY, AY, and YT

Investigation: OY, AY, YS, MK, TI, TY, SK, and GO

Visualization: OY

Funding acquisition: YO, KK, HT, MK, and OY

Data curation: AY, YS, YO, GO, MT, MN, TI, TH, YY, TM, TO, HT, HT, JN, HT, KH, KK, NO, OA, HI, TY, SK, and SCT

Project administration: OY, YO, TH, KK, HI, TY, SK, SCT, and MK

Supervision: OY

Writing–original draft: OY and MK

Writing–review & editing: All authors

