## Supplemental Materials for "Computational Mechanisms of Neuroimaging Biomarkers Uncovered by Multicenter Resting-State fMRI Connectivity Variation Profile"

### **Supplementary Table 1.** Signal-to-noise ratio (SNR) estimates of the FCs and the LASSO classifier outputs for MDD, ASD and SCZ

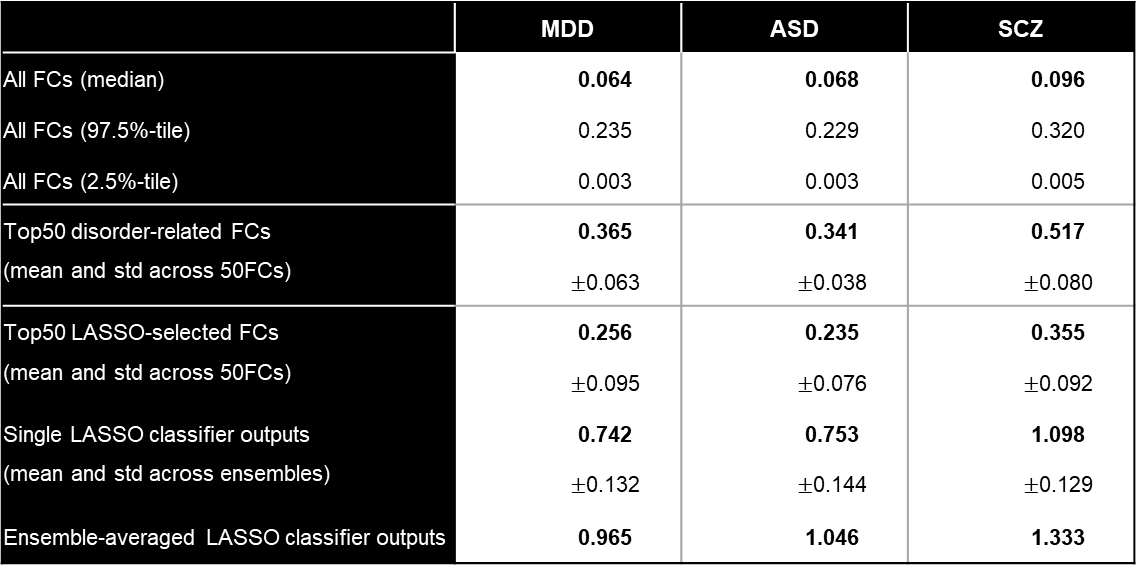

### **Supplementary Table 2.** Summary of the functional magnetic resonance imaging (fMRI) protocols

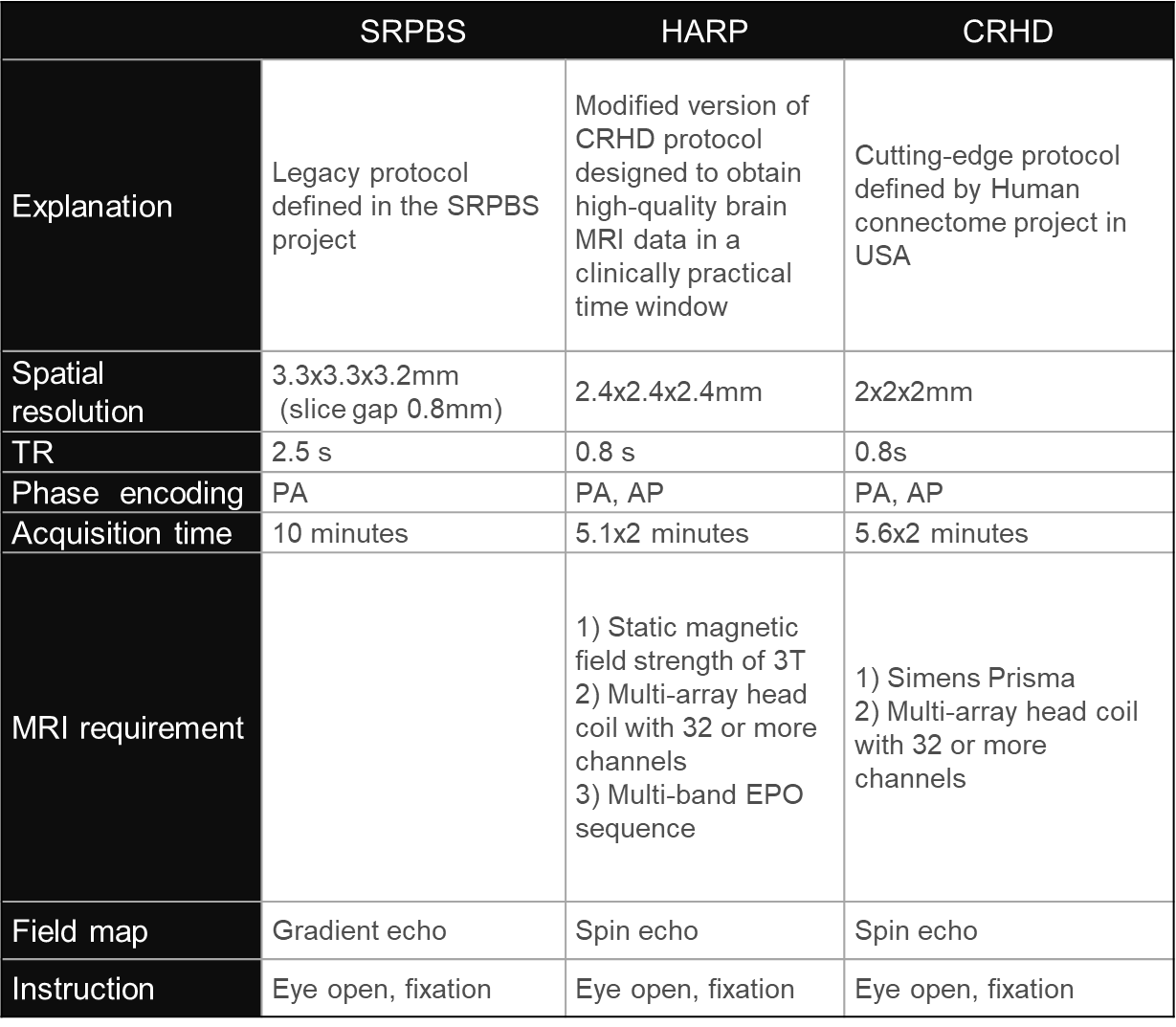

### **Supplementary Table 3.** Summary of the Brain/Minds Beyond (BMB) traveling-subject dataset

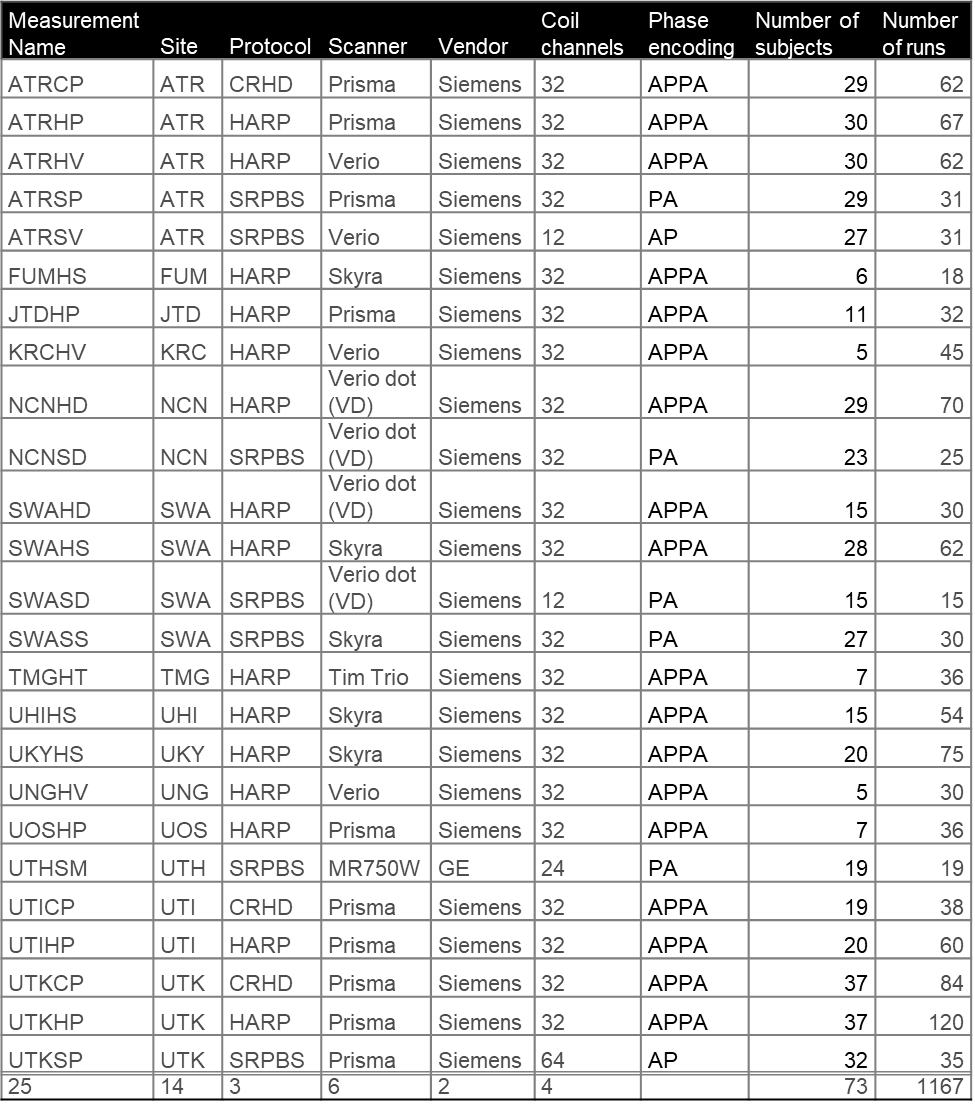

### **Supplementary Table 4.** Summary of the Strategic Research Program for Brain Sciences (SRPBS) traveling-subject dataset

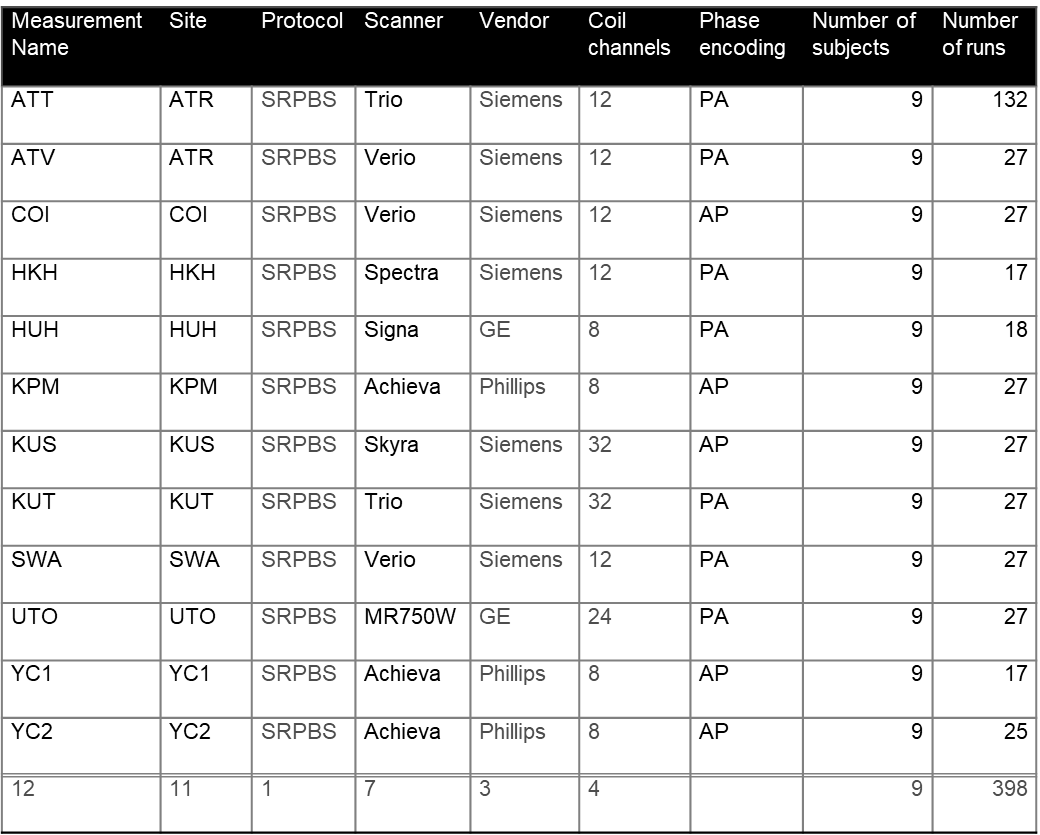

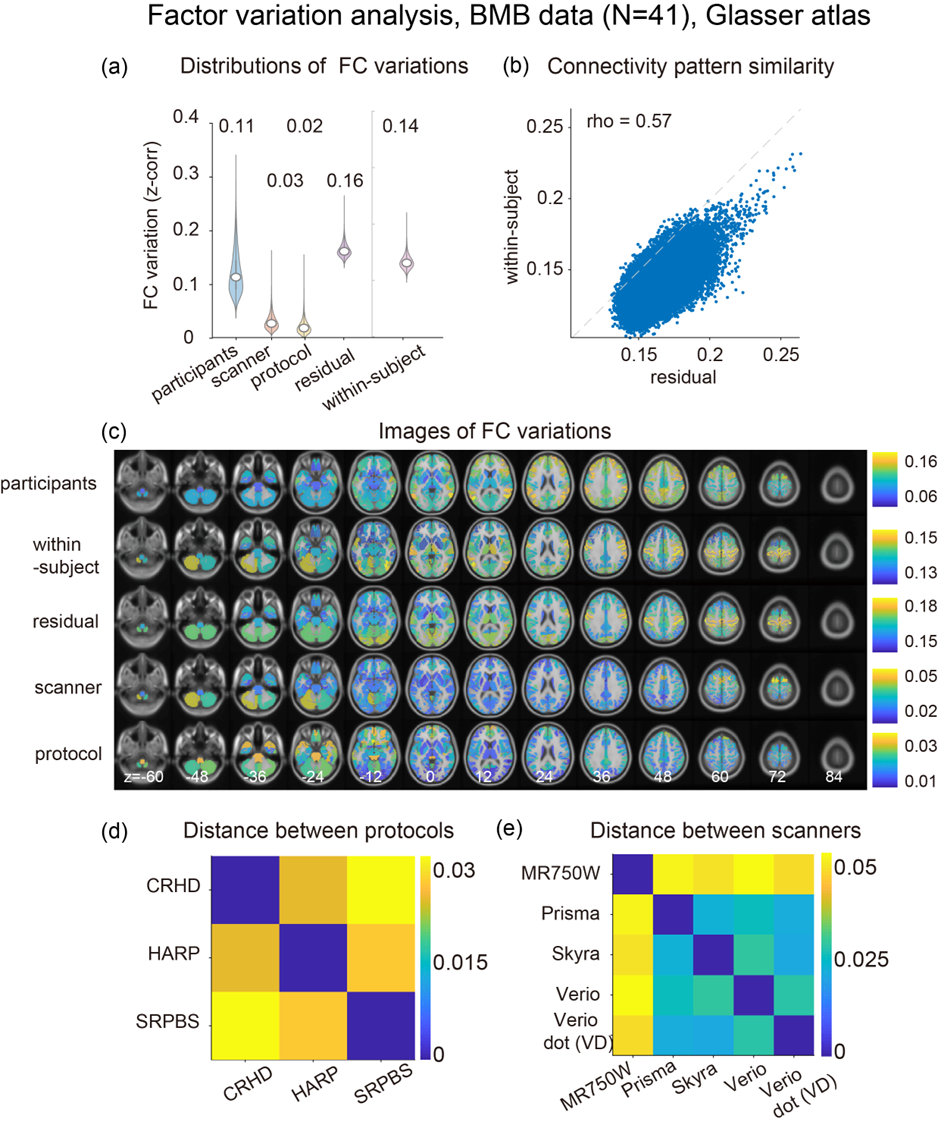

### **Supplementary Figure 1.** Summary of the factor variation analysis conducted with a non-overlapping data separation approach for the Brain Mind Beyond (BMB) dataset based on Glasser’s Multimodal Parcellation (MMP) atlas.

(a) Magnitudes of the functional connectivity (FC) variations related to each factor. (b) Comparison of connectivity pattern similarity between the residual component and within-subject variations. (c) Brain mapping of the FC variations for each factor. (d)(e) Pair-wise distances between the members of the protocol and scanner factors, respectively. All reported values are represented by z-transformed Pearson correlation coefficients.

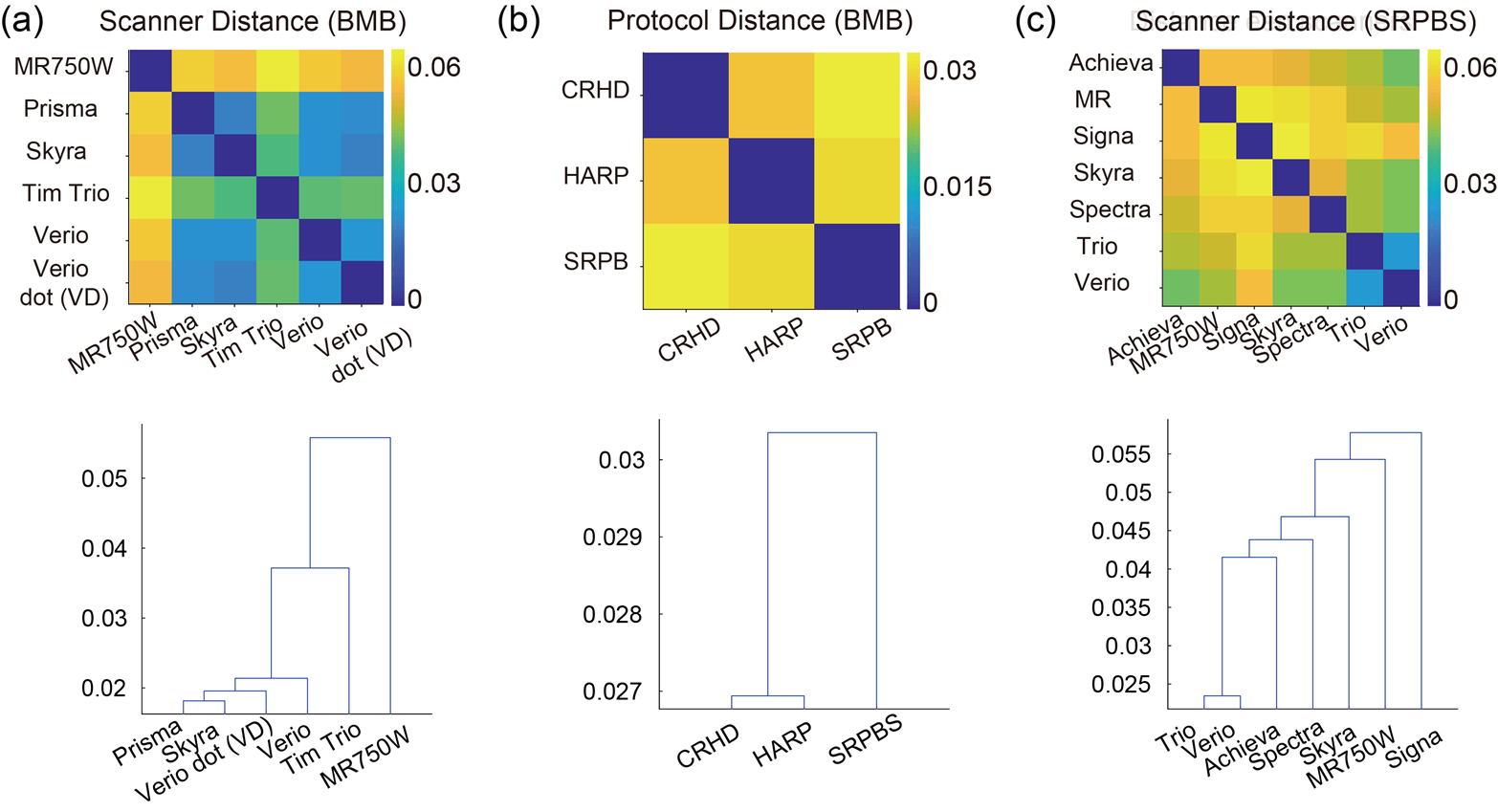

### **Supplementary Figure 2.** Distance matrices and dendrograms for the scanner and imaging protocol factors.

The distance between two protocols or two scanners was defined as the mean absolute difference between the corresponding estimated parameter vectors across all connections, and the data are represented as either a matrix (upper panels) or a dendrogram obtained by applying the agglomerative hierarchical clustering to the matrix (lower panels). (a) Scanner distance for the Brain/Minds Beyond (BMB) data. (b) Imaging protocol distance for the BMB data. (c) Scanner distance for the Strategic Research Program for Brain Science (SRPBS)  data. It is notable that the distance matrices are symmetric. The reported values are represented as z-transformed Pearson correlation coefficients. CRHD, Connectomes Related to Human Diseases; HARP, Harmonized Protocol.

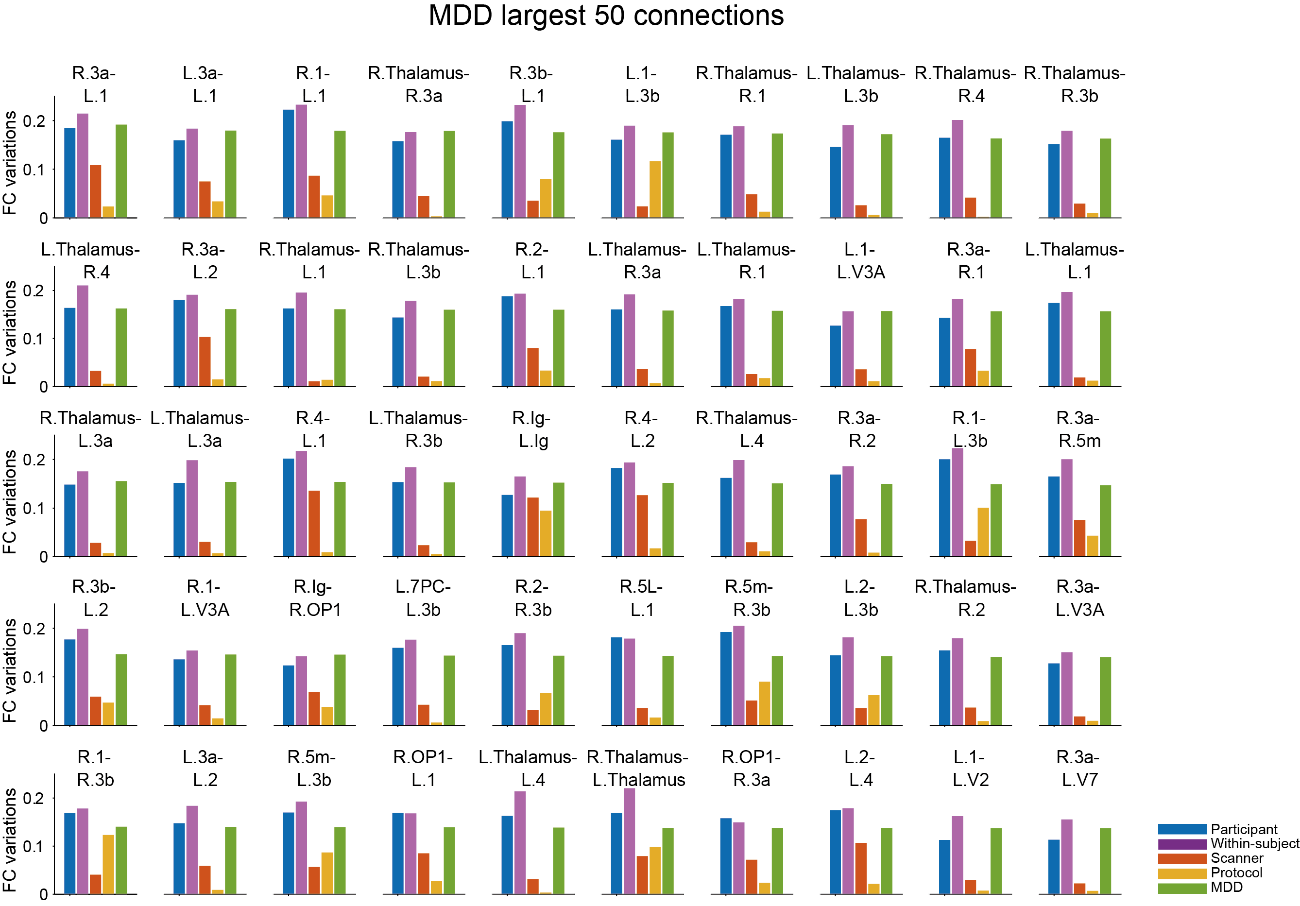

### **Supplementary Figure 3.** Functional connectivity (FC) variations at the 50 largest major depressive disorder (MDD)-related connections**.**

Each panel displays the FC variations attributed to the participant, within-subject, scanner, imaging protocol, and MDD factors for a specific connectivity. The connections are arranged in order, with the MDD-related FC differences decreasing from the top-left to the bottom-right panels. Regions are labelled in accordance with Glasser et al., 2016^1^. The FC variations are represented as z-transformed Pearson correlation coefficients. The 50 connections presented here are identical to those in Fig. 4(d).

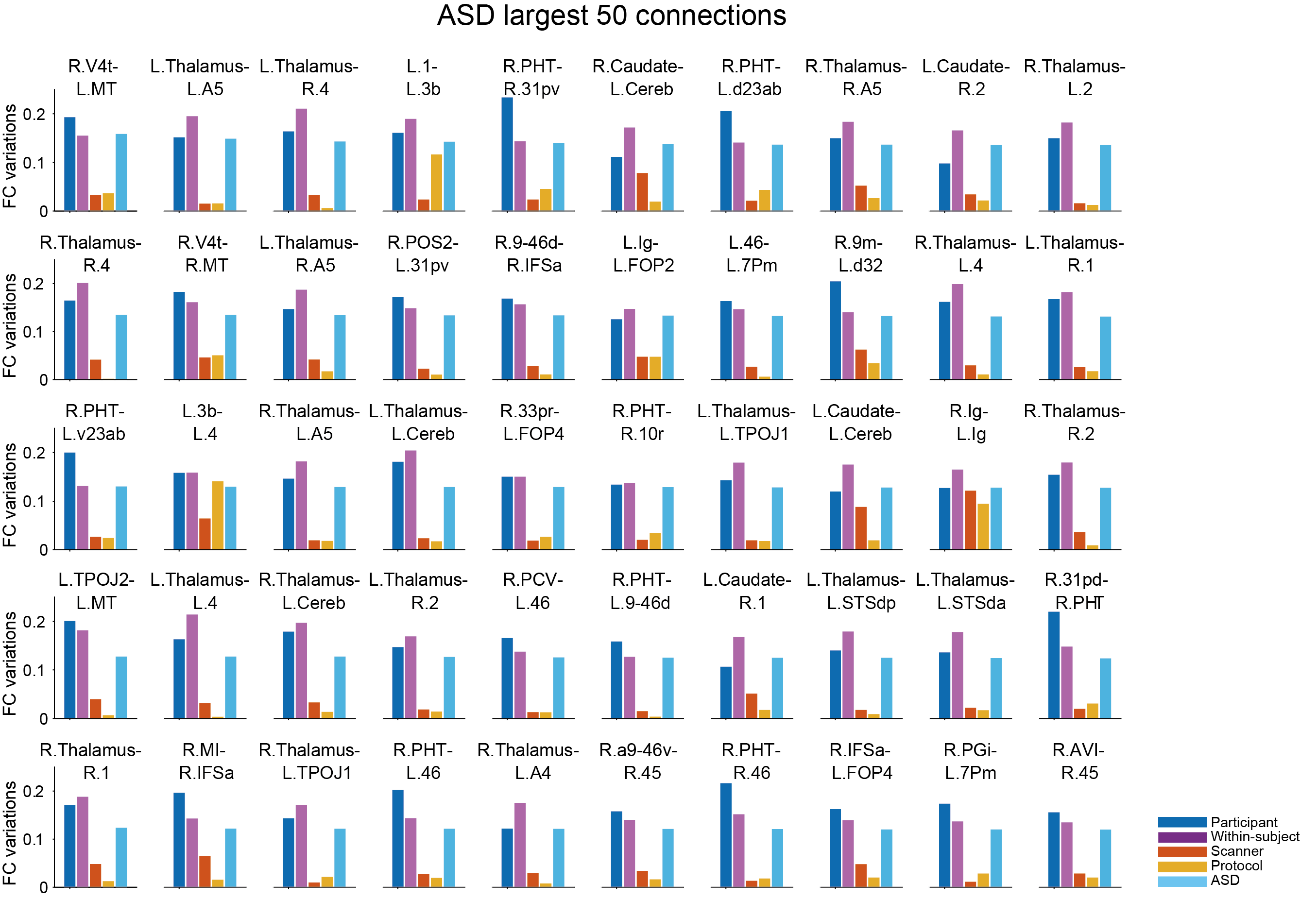

### **Supplementary Figure 4.** Functional connectivity (FC) variations at the 50 largest autism spectrum disorder (ASD)-related connections.

Each panel displays the FC variations attributed to participant, within-subject, scanner, imaging protocol, and ASD-related factors for a specific connectivity. The connections are arranged in order, with the ASD-related FC variations decreasing from the top-left to the bottom-right panels. Regions are labelled in accordance with Glasser et al., 2016. The FC variations are represented as z-transformed Pearson correlation coefficients. The 50 connections presented here are identical to those in Fig. 4(d).

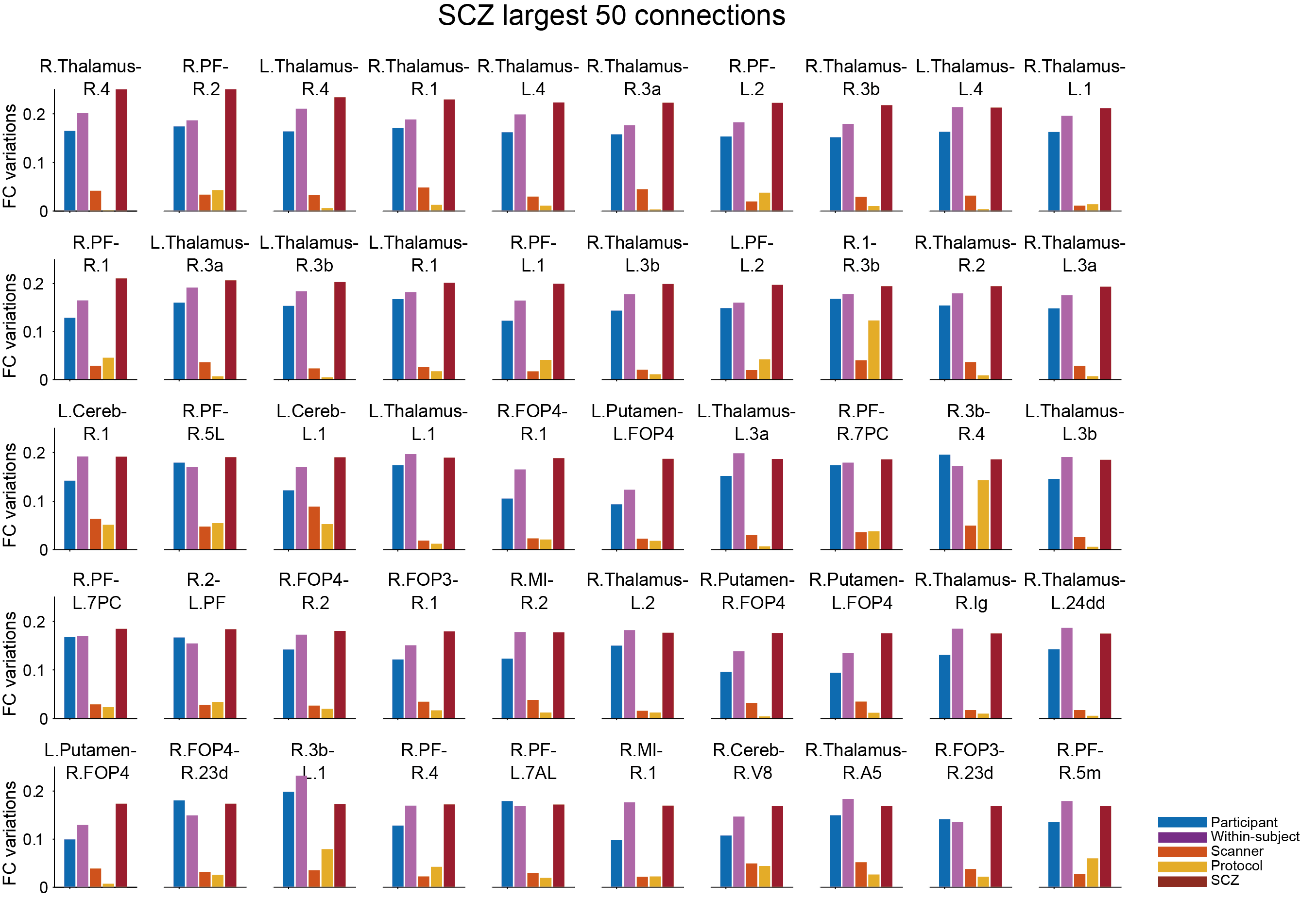

### **Supplementary Figure 5.** Functional connectivity (FC) variations at the 50 largest autism spectrum disorder (SCZ)-related connections.

Each panel displays the FC variations attributed to participant, within-subject, scanner, imaging protocol, and SCZ-related factors for a specific connectivity. The connections are arranged in order, with the SCZ-related FC variations decreasing from the top-left to the bottom-right panels. Regions are labelled in accordance with Glasser et al., 2016. The FC variations are represented as z-transformed Pearson correlation coefficients. The 50 connections presented here are identical to those in Fig. 4(d).

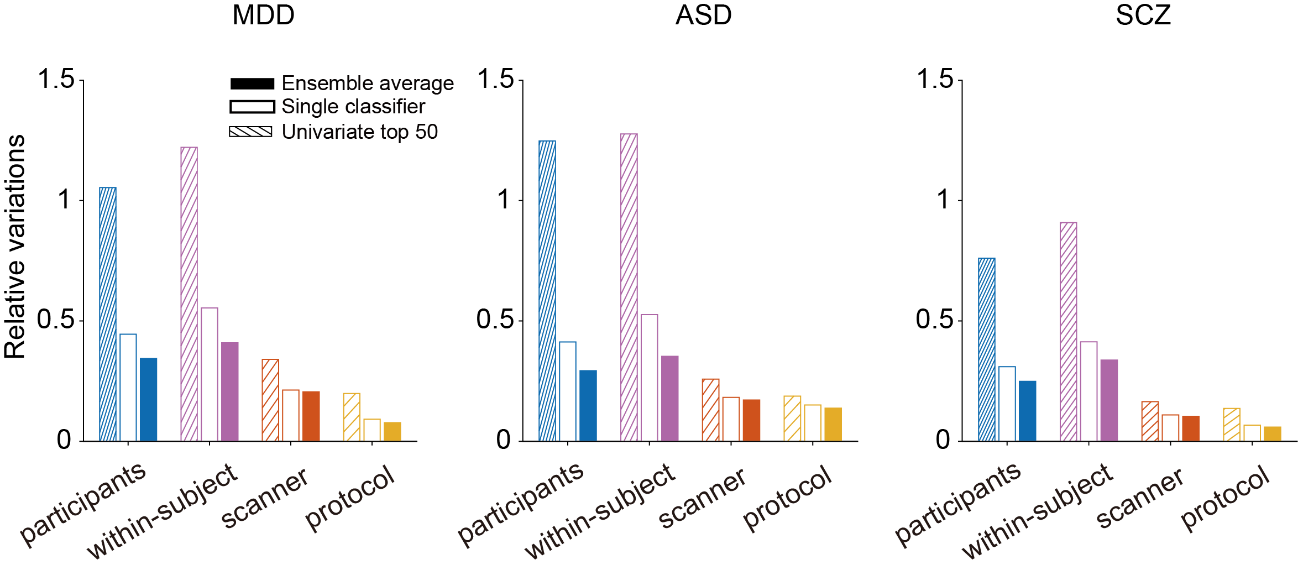

### **Supplementary Figure 6.** Comparison of relative variations for each psychiatric disorder.

The variations due to participant, within-subject across-runs, scanner, and protocol factors relative to that of the disease factor are compared for the following three outputs: the average of the top 50 largest disorder-related FCs, the multivariate FC biomarker output before ensemble averaging, and the multivariate FC biomarker output after ensemble averaging. The relative variations were computed by dividing each factor variation by the disease difference for major depressive disorder (MDD), autism spectrum disorder (ASD), and schizophrenia (SCZ) from left-to-right, respectively. Compared with the impact based on simple averaging of the top 50 largest disorder-related FCs, the use of multivariate FC biomarkers reduced the variations due to disorder-unrelated factors (participant, within-subject across-runs, scanner, and protocol difference) by more than half.

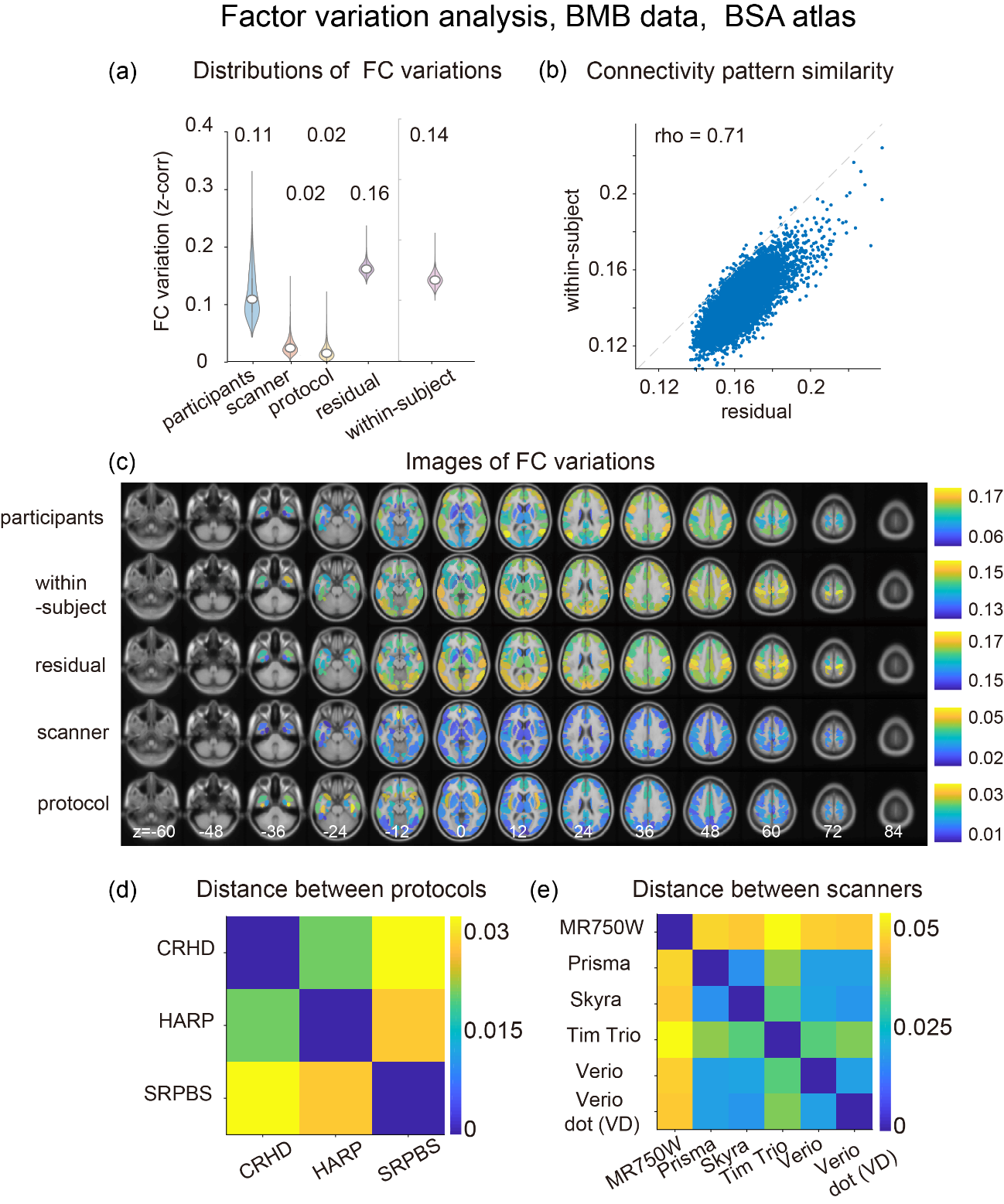

### **Supplementary Figure 7.** Summary of the functional connectivity (FC) variation analysis for the Brian/Mind Beyond (BMB) data conducted using the coarse anatomical brain atlas (BrainVISA Sulci Atlas).

A total of 137 regions were included in the analysis, excluding the cerebellar areas. (a) Distributions of the magnitudes of the FC variations. (b) Comparison of connectivity pattern similarity between the residual component and within-subject variations. (c) Brain mapping of FC variations. (d)(e) Pair-wise distances between members of the imaging protocol and scanner factors, respectively . All reported values are represented by z-transformed Pearson correlation coefficients. CRHD, Connectomes Related to Human Diseases; HARP, Harmonized Protocol; SRPBS , Strategic Research Program for Brain Sciences. The BrainVisa Sulci atlas can be obtained from https://brainvisa.info/web/morphologist.html.

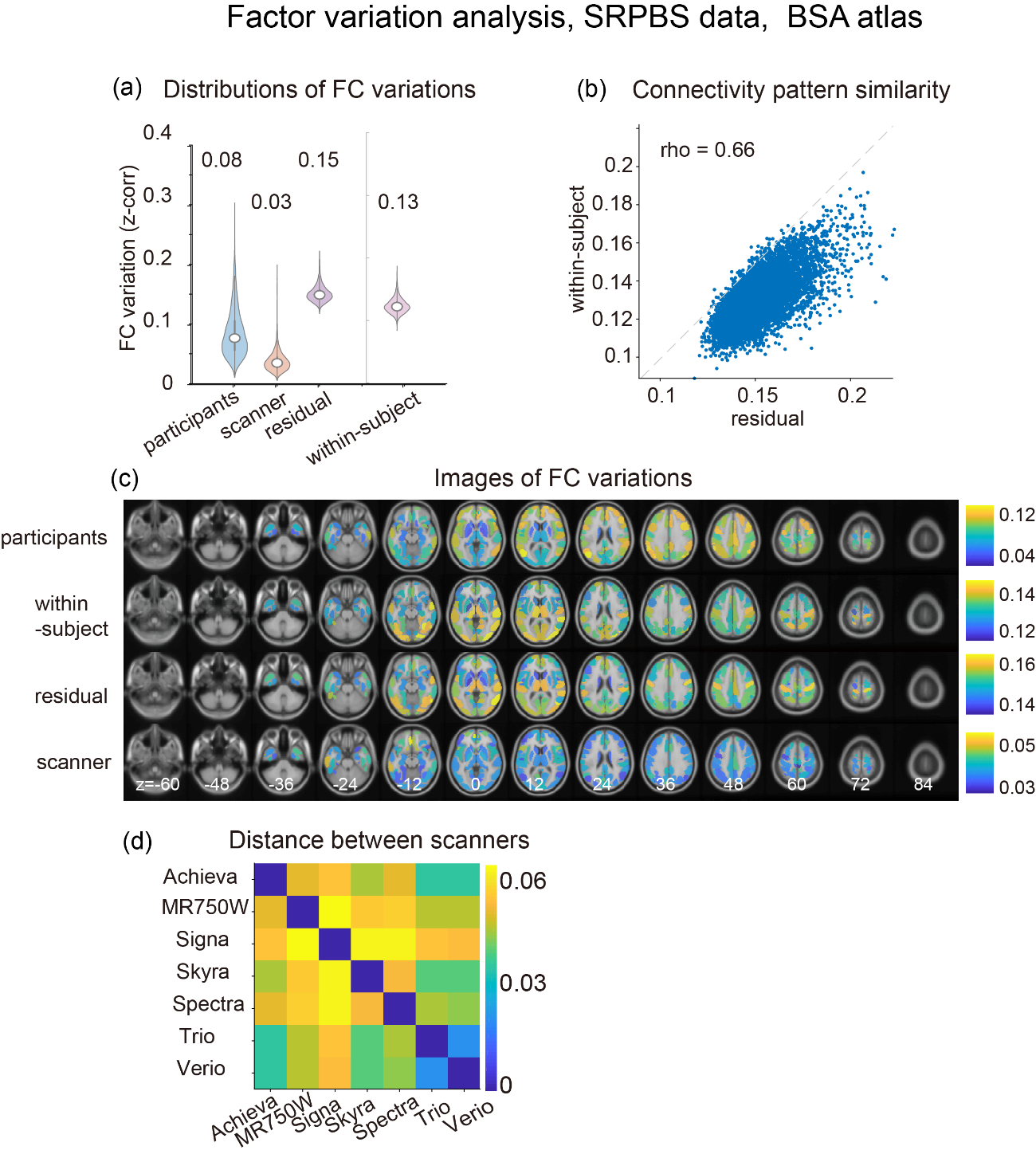

### **Supplementary Fig. 8.** Summary of the functional connectivity (FC) variation analysis for the Strategic Research Program for Brain Sciences (SRPBS) dataset conducted using the coarse anatomical brain atlas (BrainVISA Sulci Atlas).

A total of 137 regions were included in the analysis, with the exclusion of the cerebellar areas. Each panel retains an identical format to the panels depicted in Supplementary Fig. 7.

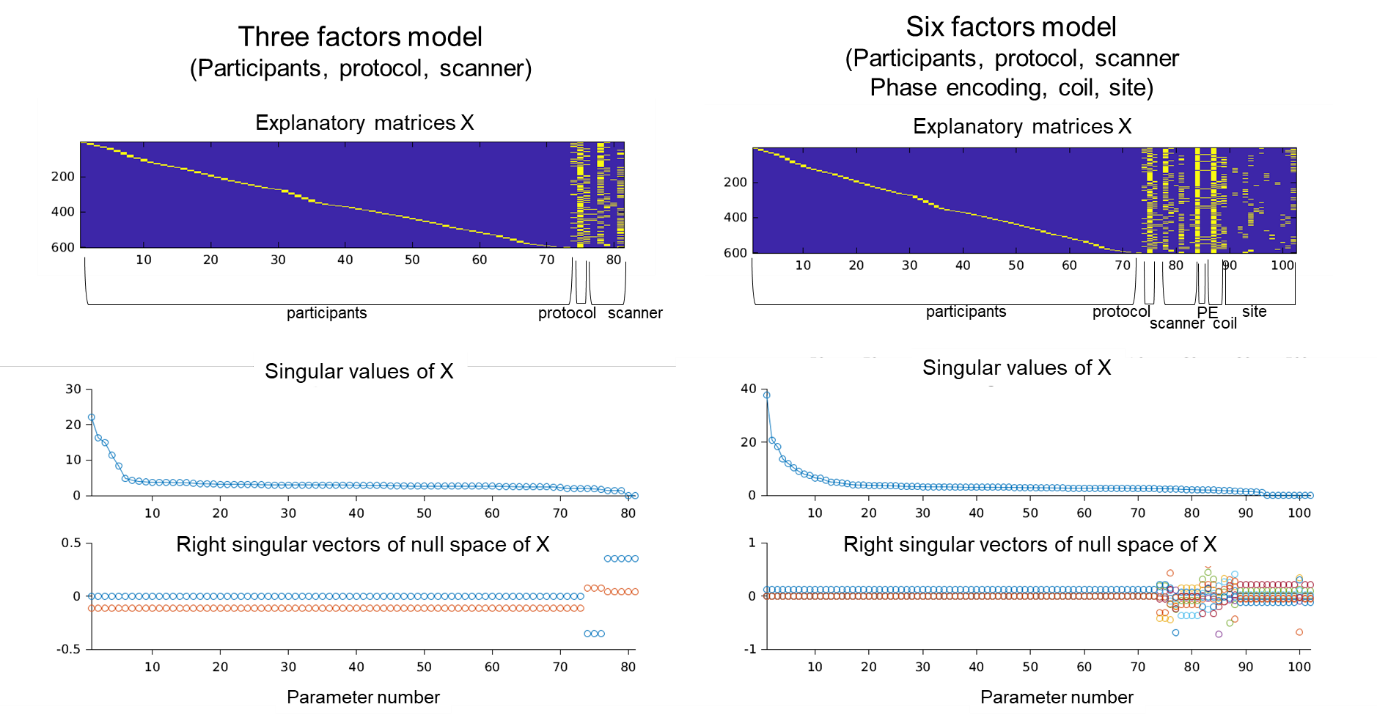

### **Supplementary Figure 9.** Model checking using the singular value decomposition (SVD) of the explanatory matrix.

From top to bottom, the explanatory matrix, singular values, and null space are displayed, respectively, for three-factor models consisting of the participant, protocol, and scanner factors (left panels) and six-factor models consisting of the participant, protocol, scanner, phase encoding, coil, and site factors (right panels).

We consider the case where there are more than or equal to two factors of interest and each factor variable is the categorical variable. In this case, the parameters of the linear fixed effect model has components which cannot be determined by the data. Thus it is important to confirm what is undetermined for further analysis of the estimated parameters. The singular value decomposition (SVD) of the explanatory matrix gives a way to understand the undetermined components.
 Let $\boldsymbol{X}$ and $\boldsymbol{\beta}$ denote the binary explanatory matrix and the parameter vector consisting of the factors of interest, respectively. We further assume the number of data is larger than the number of parameters. To factorize the magnitude of each factor on the functional connectivity $\boldsymbol{z}$, we assumed the linear regression model,
$\boldsymbol{z}=\boldsymbol{X\beta+}\boldsymbol{\epsilon}$**.** …(s1)
Minimizing the square errors leads to the normal equation which the parameter vector $\boldsymbol{\beta}$ must meet,
$\boldsymbol{X}^{\boldsymbol{t}}\boldsymbol{X\beta=}\boldsymbol{X}^{\boldsymbol{t}}\boldsymbol{z}$**.** …(s2)
 As the explanatory matrix $\boldsymbol{X}$ is not of column full rank (easily confirmed by summing the columns within each, the inverse of $\boldsymbol{X}^{\boldsymbol{t}}\boldsymbol{X}$ does not exist. Instead the parameter vector $\boldsymbol{\beta}$ is obtained using the Moore-Penrose generalized inverse matrix,
${\boldsymbol{\beta=}{\left( \boldsymbol{X}^{\boldsymbol{t}}\boldsymbol{X} \right)^{\boldsymbol{+}}\boldsymbol{X}}^{\boldsymbol{t}}\boldsymbol{z}}$. …(s3)
 This solution has the minimum L2 norm among the possible solutions which meet the normal equation. The SVD of the explanatory matrix provides more insight to this solution. As the explanatory matrix is not of column full rank, there exist at least one zero singular values. Let $\boldsymbol{s}_{\boldsymbol{+}}$ denote a diagonal matrix whose diagonal elements are the positive singular values and $V_{+}$ and $V_{0}$ denote the right singular vectors corresponding to the positive singular values and zero singular values, respectively. Then the SVD of the explanatory matrix is written by

$$\boldsymbol{X}\boldsymbol{＝}\boldsymbol{U}\left[ \begin{matrix} \boldsymbol{s}_{\boldsymbol{+}} & \boldsymbol{0} \\ \boldsymbol{0} & \boldsymbol{0} \end{matrix} \right]\left[ \begin{matrix} \boldsymbol{V}_{\boldsymbol{+}} & \boldsymbol{V}_{\boldsymbol{0}} \end{matrix} \right]^{\boldsymbol{t}}\boldsymbol{,}$$

By substituting this equation to equation (s3), we have

$\boldsymbol{\beta=}{\left[ \begin{matrix} \boldsymbol{V}_{\boldsymbol{+}}\boldsymbol{s}_{\boldsymbol{+}}^{\boldsymbol{-1}} & \boldsymbol{V}_{\boldsymbol{0}}\boldsymbol{0} \end{matrix} \right]\boldsymbol{U}}^{\boldsymbol{t}}\boldsymbol{z}$***.*** …(s4)
Here we used the generalized inverse of the diagonal matrix given by
$\left[ \begin{matrix} \boldsymbol{s}_{\boldsymbol{+}}^{\boldsymbol{2}} & \boldsymbol{0} \\ \boldsymbol{0} & \boldsymbol{0} \end{matrix} \right]^{\boldsymbol{+}}\boldsymbol{=}\left[ \begin{matrix} \boldsymbol{s}_{\boldsymbol{+}}^{\boldsymbol{-2}} & \boldsymbol{0} \\ \boldsymbol{0} & \boldsymbol{0} \end{matrix} \right]$**.**
 From equation (s4), we can know the estimated parameter does not have any information on the space spanned by $V_{0}$. The space spanned by $V_{0}$ is called the null space. Visualizing the singular values and the null space provides the information about what is undetermined in the estimated parameter.

Supplementary Figure 9 presents the explanatory matrix, singular values, and the null space for two types of factor models. The first type includes three factors, namely participants, protocol, and scanner factors. In this case, there are two zero singular values, and their corresponding right singular vectors display a constant value within participants, protocol, and scanner factors. This indicates that determining the baseline value of each factor is not feasible. The second type of factor model consists of six factors: participants, protocol, scanner, phase encoding, coil, and site. For this model, nine zero singular values were observed, and the corresponding right singular vectors displayed complex patterns due to co-linearity between the factors. Consequently, it is difficult to investigate the estimated parameters for further analysis.

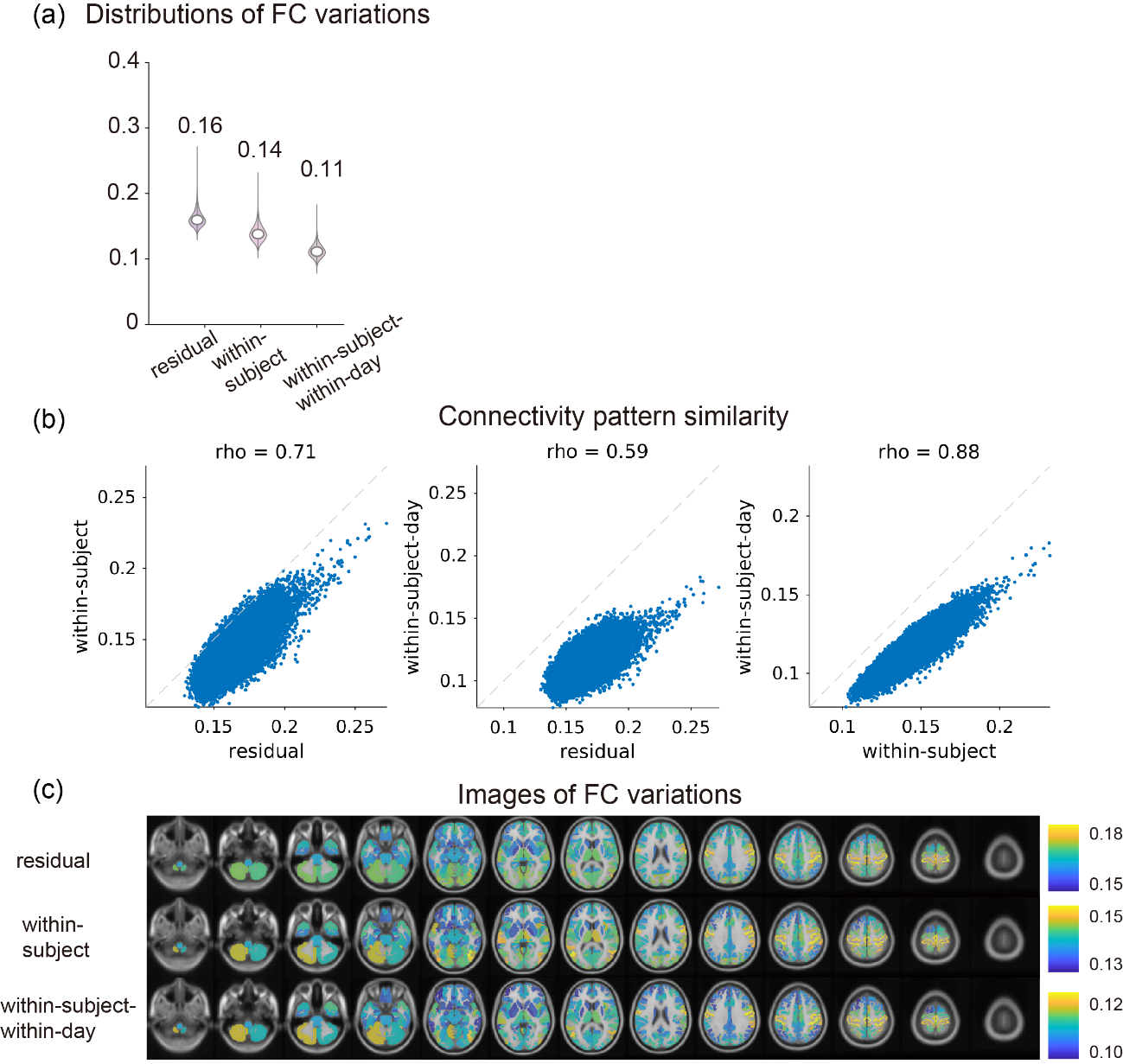

### **Supplementary Fig. 10.** Analysis of within-subject functional connectivity (FC) variation and within-subject within-day FC variation for the Brian/Minds Beyond (BMB) traveling-subject dataset based on the Glasser Multimodal Parcellation (MMP) atlas.

The within-subject FC variation analysis described in the main text included the between-days variations, as each subject’s data was collected from at least two days of experimental sessions, with three runs per day. To consider the within-subject FC variation in a single day, the subject-and-day-specific FC pattern, which was obtained by averaging results over three runs in one day, was subtracted from the data from each run conducted on the same day to calculate the within-subject within-day FC variations. Subsequently, the within-subject within-day FC variation was obtained as the standard deviation of the within-subject within-day FC deviations collected from all days. (a) Magnitude distributions of the residual component, within-subject, and the within-subject within-day FC variation. (b) Comparison of connectivity pattern similarity. (c) Brain mapping of FC variations.

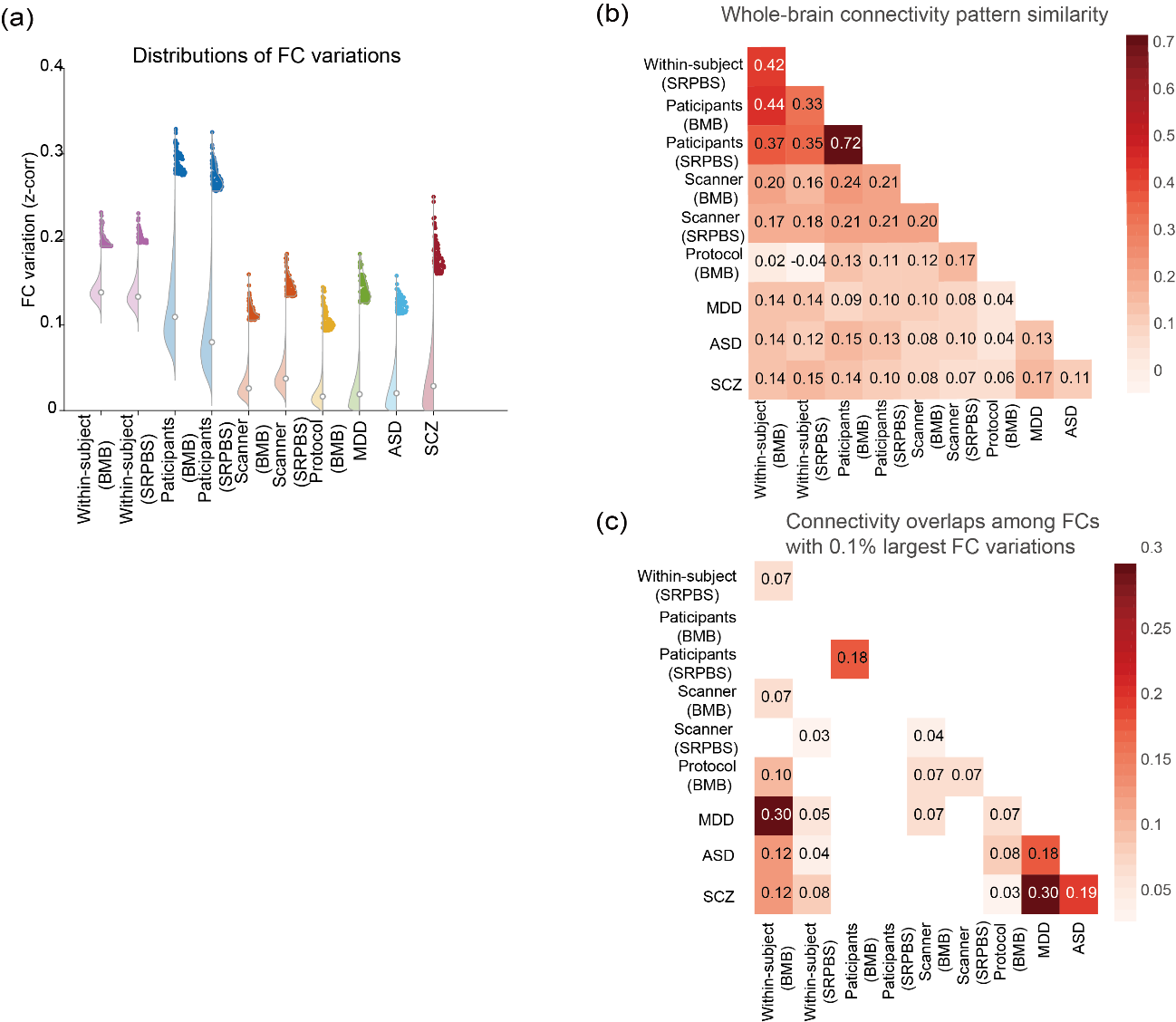

### **Supplementary Figure 11.** Summary of the functional connectivity (FC) variation analysis.

(a) The left-half of each violin plot depicts the distribution of FC variations for the whole-brain connectivity (71,631 connections), and the right-half of each violin plot depicts the distribution of the FC variation for the connections with the 0.1% largest FC variation for each factor (72 connections). (b) Comparison of whole-brain connectivity pattern similarity between all pairs of the factors, as quantified by the rank correlation analysis. (c) Overlapping connectivity among the connections with the 0.1% largest FC variations. For each pair of factors, the consistency of the selected connectivity was quantified by the Sorensen-Dice coefficient . Only the statistically significant pairs (p<0.05, with Bonferroni correction) are shown.

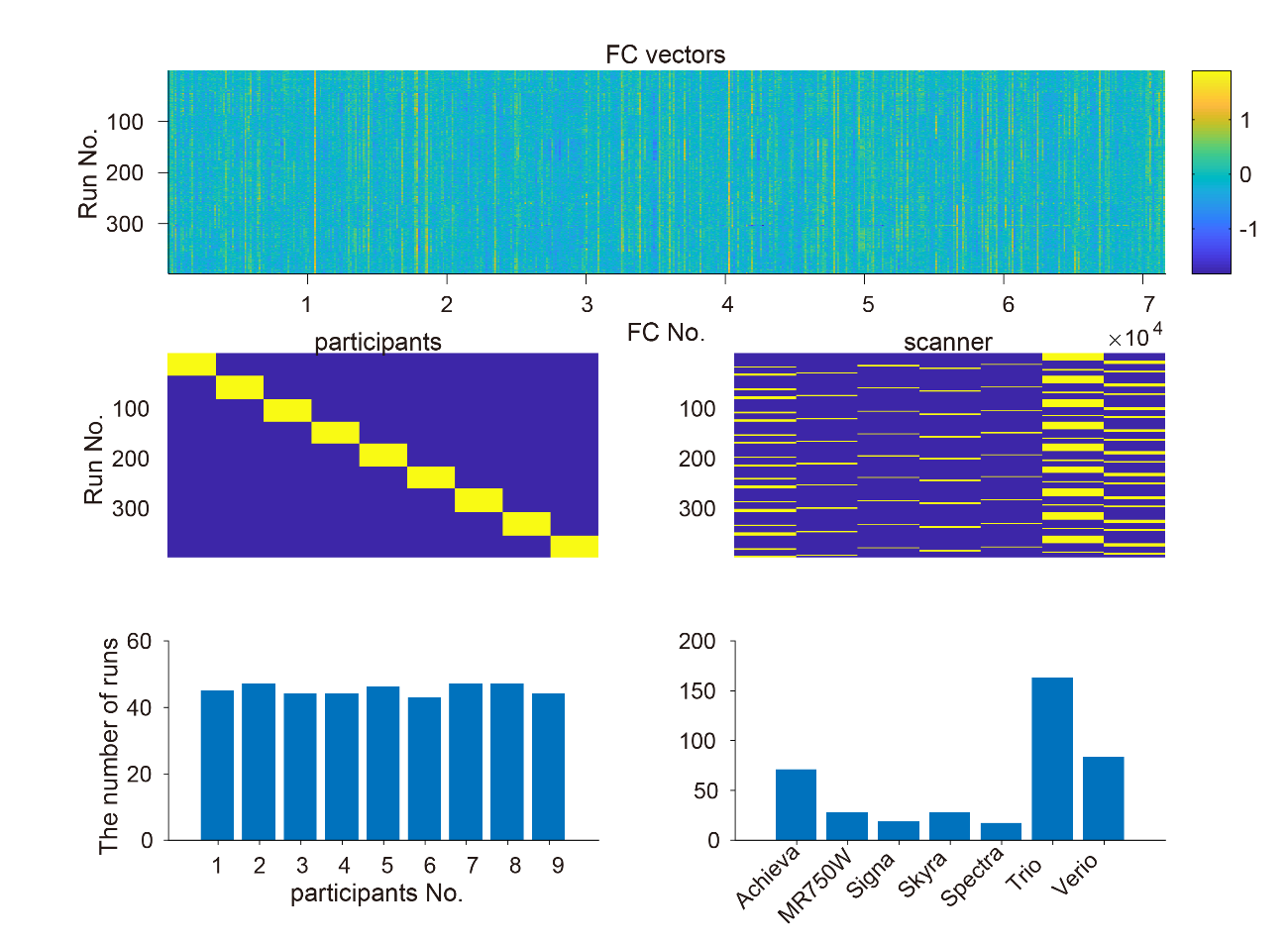

### **Supplementary Figure 12.** Graphical summary of the Strategic Research Program for Brain Sciences (SRPBS) traveling-subject data used for analysis (398 runs, nine subjects, and seven scanners).

The top section illustrates the FC vectors of all 398 runs computed via Glasser’s multimodal parcellation for 379 brain regions. Each row vector represents a vectorized whole-brain FC  (379*378/2 = 71,631 connections) derived from a 10-minute eyes-open resting-state experiment. Each element of the matrix is a Fisher’s z-transformed Pearson correlation coefficient. In the middle row, from left to right, one-hot encoding matrices of the participants (subjects) and scanners are presented, respectively, which were used as the explanatory variables in the linear fixed effects model. The graphs in the bottom row present the total number of runs plotted against each member of the participant and scanner factors, respectively.

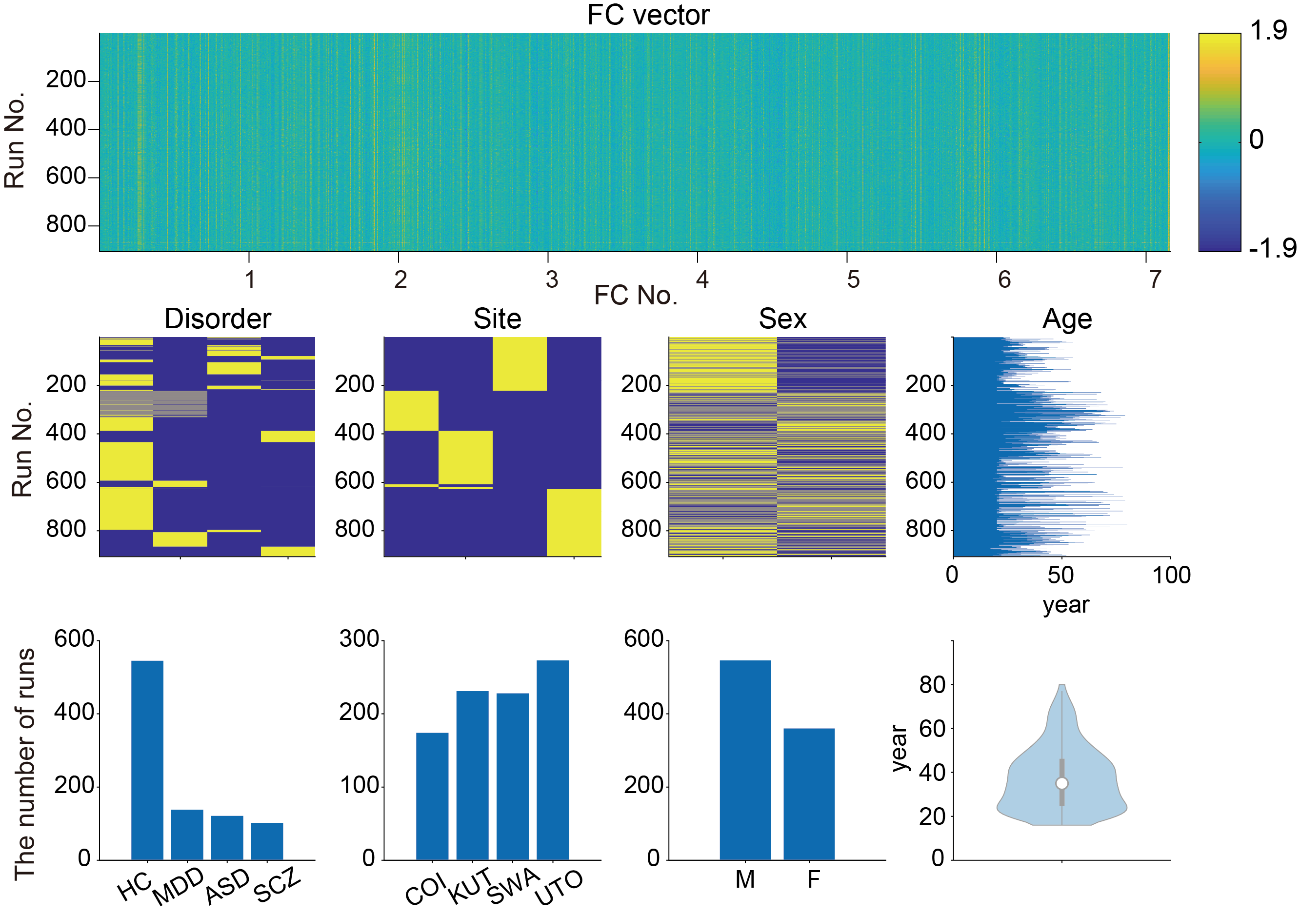

### **Supplementary Figure 13.** Graphical summary of the Strategic Research Program for Brain Sciences (SRPBS) multi-disorder data used for analysis (906 subjects runs from four sites).

The top section illustrates the functional connectivity (FC) vectors of all the runs computed using Glasser’s multimodal parcellation for 379 brain regions. Each row  vector represents a vectorized whole-brain FC  (379*378/2 = 71,631 connections) derived from a 10-minute eyes-open resting-state experiment. Each element of the matrix is a Fisher’s z-transformed Pearson correlation coefficient. In the middle row, from left to right, are the one-hot encoding matrices for the psychiatric disorders, sites, and sex, followed by a bar graph representing the number of runs plotted against the subject ages, respectively. The graphs in the bottom row presents the summary statistics of the number of runs for each psychiatric disorder, site, sex, and age, respectively. HC, healthy control; MDD, major depressive disorder; ASD, autism spectrum disorder; SCZ, schizophrenia; COI, Center of Innovation at Hiroshima University ; KUT, Kyoto University ; SWA, Showa University; UTO, University of Tokyo;  M, male; F, female.

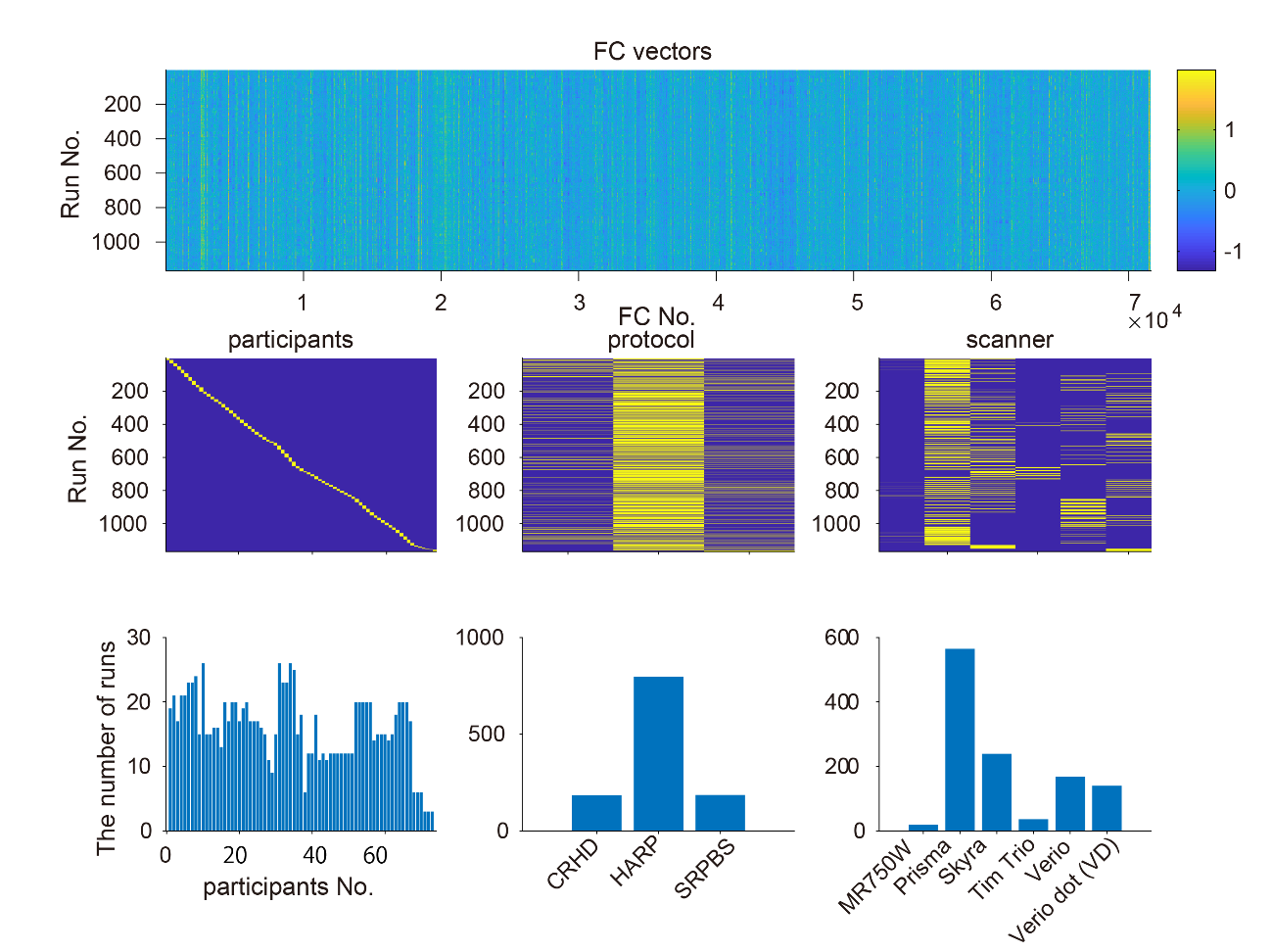

### **Supplementary Figure 14.** Graphical summary of the Brain/Minds Beyond (BMB) traveling-subject data used for analysis (1,167 runs, 73 subjects, three protocols, and six scanners).

The top section illustrates the functional connectivity (FC) vectors of all 1,167 runs computed via Glasser’s multimodal parcellation for 379 brain regions. Each row vector represents a vectorized whole-brain FC  (379*378/2 = 71,631 connections) derived from a 10-minute eyes-open resting-state experiment. Each element of the matrix is represented as a Fisher’s z-transformed Pearson correlation coefficient. In the middle section, from left to right, one-hot encoding matrices for participants (subjects), protocols, and scanners are presented, respectively. These matrices were used as the explanatory variables in the linear fixed effects modeling. The bottom section presents the total number of runs corresponding to each member of the participant, protocol, and scanner factors, respectively. CRHD, Connectomes Related to Human Diseases; HARP, Harmonized Protocol; SRPBS , Strategic Research Program for Brain Sciences.
